# Assessing donor-to-donor variability in human intestinal organoid cultures

**DOI:** 10.1101/2021.07.23.453590

**Authors:** Sina Mohammadi, Carolina Morell-Perez, Charles W. Wright, Thomas P. Wyche, Cory H. White, Theodore R. Sana, Linda A. Lieberman

**Author notes:** Current address: Health and Environmental Sciences Institute, Washington, DC 20005 USA.

## Abstract

Donor-to-donor variability in primary human organoid cultures has not been well characterized. As these cultures contain multiple cell types, there is greater concern that variability could lead to increased noise. In this work we investigated donor-to-donor variability in human gut adult stem cell (ASC) organoids. We examined intestinal developmental pathways during culture differentiation in ileum- and colon-derived cultures established from multiple donors, showing that differentiation patterns were consistent among cultures. This finding indicates that donor-to-donor variability in this system remains at a manageable level. Intestinal metabolic activity was evaluated by targeted analysis of central carbon metabolites and by analyzing hormone production patterns. Both experiments demonstrated similar metabolic functions among donors. Importantly, this activity reflected intestinal biology, indicating that these ASC organoid cultures are appropriate for studying metabolic processes. This work establishes a framework for generating high confidence data using human primary cultures through thorough characterization of variability.

## Introduction

Historically, intestinal biology has been studied using immortalized or transformed cell lines. While these cell lines play a useful role, primary cell culture models provide a more physiologically relevant system to address unanswered questions and more accurately model *in vivo* responses. Groundbreaking work in 2009 described modern organotypic cultures that are derived from adult mouse intestinal stem cells, are self-organizing, and contain nearly a complete repertoire of functional (terminally differentiated) cells as found in the originating tissue (Sato et al., 2009). A system derived from human intestinal stem cell was described subsequently (Sato et al., 2011a). These adult stem cell (ASC) organoid cultures enable a vast swath of *in vitro* studies in cells derived from a healthy host, in contrast to previous culture systems that relied on immortalized or transformed cells.

Gut ASC organoid cultures containing differentiated cells recapitulate *in vivo* intestinal cell composition. Absorptive and secretory cell types, which carry out the function of the intestine, are present in ASC organoid cultures. Enterocytes serve as absorptive cells in the intestine and are the most abundant cell type (Fish and Burns, 2020). Their function is quite diverse, ranging from lipid, sugar, amino acid, and inorganic molecule metabolism to bile acid reabsorption (Chen et al., 2018; Dawson, 2011; Gao et al., 2019; Hernando and Wagner, 2018; Hussain, 2014; Ko et al., 2020). Three secretory cell types commonly found in the intestine are also found in ASC organoids. Paneth cells are specialized secretory cells that reside at the base of intestinal crypts. In addition to secreting antimicrobial peptides and proteins, Paneth cells produce essential niche signals required for stem cell maintenance (Clevers and Bevins, 2013). Goblet cells produce and secrete mucus (Johansson et al., 2013), thereby generating a barrier surface. Enteroendocrine cells produce and secrete neurotransmitters and incretin hormones (Gribble and Reimann, 2019), that play major roles in gut peristalsis and vasodilation as well as pancreatic insulin secretion and satiety. Two other cell types (tuft cells (Nevo et al., 2019) and microfold (M) cells (Dillon and Lo, 2019)) occur *in vivo* but differentiation towards these lineages has to be induced by the addition of cytokines in ASC organoid cultures.

As ASC organoids contain both stem and differentiated cells, they have been deployed in a range of studies including fundamental pathway analysis efforts (Date and Sato, 2015; Dubey et al., 2020; Lindemans et al., 2015), establishing disease models (Bartfeld, 2016; Schutgens and Clevers, 2020; van der Vaart and Clevers, 2020), validating drug activity and efficacy, and genetic screening efforts (Michels et al., 2020; Ringel et al., 2020). Despite wide deployment, to date, variations among donors in differentiation patterns and therefore functional readouts have not been characterized in human ASC intestinal organoid cultures. To address this knowledge gap, we established cultures from ileum and colon of six donors and examined donor-to-donor variability for multiple parameters. We show a high degree of correlation among donors, indicating that differentiation patterns are consistent. Additionally, small differences among donors in fundamental intestinal functions (such as metabolic activity and hormone secretion) did not hinder interpretation of results. Taken together these data highlight the importance of thoroughly characterizing human ASC organoid cultures with a diverse set of assays. Heterogeneity is an inherent and appropriate feature of human ASC organoid cultures but we show that variability could be managed to yield robust and interpretable datasets.

## Results

### Assessing developmental gene expression patterns

Gut adult stem cell (ASC) organoid cultures were established by isolating crypts from human intestinal tissue as previously described (Miyoshi and Stappenbeck, 2013; Sato et al., 2011a). Cultures were derived from ileum and/or colon of donated cadaver tissues from six adult donors of diverse ages and ethnicities (Table 1). Both ileum and colon cultures were established from all donors except donors 4 (ileum only) and 5 (colon only). Median age and body mass index were 31.5 (range: 18-47) and 27 kg/m^2^ (range: 19.7-32.9 kg/m^2^), respectively (summarized in Table 1).

**Table 1:** Donor demographics

| donor number | age | sex | ethnicity | BMI (kg/m<sup>2</sup>) | cause of death | acquisition state or territory | tissue |  |
| --- | --- | --- | --- | --- | --- | --- | --- | --- |
|  |  |  |  |  |  |  | ileum | colon |
| 1 | 47 | male | African American | 27.2 | stroke | GA | x | x |
| 2 | 43 | female | Caucasian | 29.9 | stroke | OH | x | x |
| 3 | 27 | male | Hispano | 32.9 | trauma | CA | x | x |
| 4 | 25 | male | African American | 21.3 | stroke | MD | x | -- |
| 5 | 18 | male | African American/Hispano | 19.7 | stroke | PR | -- | x |
| 6 | 29 | male | Hispano | 31.3 | anoxia | TX | x | x |

**Table 2:** Quantitative PCR markers for assessing gut development

| transcript | TaqMan ID | marker type |
| --- | --- | --- |
| MKI67 | Hs01032443_m1 | proliferative/stem cells |
| LGR5 | Hs00969422_m1 |  |
| ASCL2 | Hs00270888_s1 |  |
| AXIN2 | Hs00610344_m1 |  |
| ATOH1 | Hs00944192_s1 | secretory cells |
| MUC2 | Hs03005103_g1 |  |
| TFF3 | Hs00902278_m1 |  |
| CHGA | Hs00900370_m1 |  |
| LYZ | Hs00426232_m1 |  |
| DEFA5 | Hs00360716_m1 |  |
| DEFA6 | Hs00427001_m1 |  |
| HES1 | Hs00172878_m1 | absorptive cells |
| PDL1/CD274 | Hs00204257_m1 |  |
| BTNL3 | Hs00757802_m1 |  |
| BTNL8 | Hs01063697_m1 |  |
| KRT20 | Hs00300643_m1 |  |
| CYP3A4 | Hs00604506_m1 |  |
| P-gp/ABCB1/CD243 | Hs00184500_m1 |  |
| BCRP/ABCG2 | Hs01053790_m1 |  |
| SLC15A1 | Hs00192639_m1 |  |
| SLC10A2/ASBT | Hs01001557_m1 |  |
| PPIB | Hs00168719_m1 | endogenous control |

These adult stem cell cultures were grown in proliferation media for two days and switched to two types of differentiation media on day three. Certain growth factors were removed (ENR media (Sato et al., 2011a)) or reduced (5% LWRN media (Miyoshi and Stappenbeck, 2013)) to induce differentiation of cultures. Because of both technical and potential impact on signaling pathways, both media formulations were included (details are included in supplemental materials).

Samples were collected at days 2, 4, 7, and 10 after the initiation of ASC organoid cultures (Figure 1A). Multiple time courses (2-3 per culture spanning several passages) were carried out for each culture to assess inter- and intra-donor variability. Furthermore, isolated crypts that were used to initiate cultures were included for the comparison of ASC organoid cultures with original tissue. Overall, 26 time-course assays were carried out, generating 556 RNA samples, leading to 13344 quantitative PCR observations.

**Figure 1:**
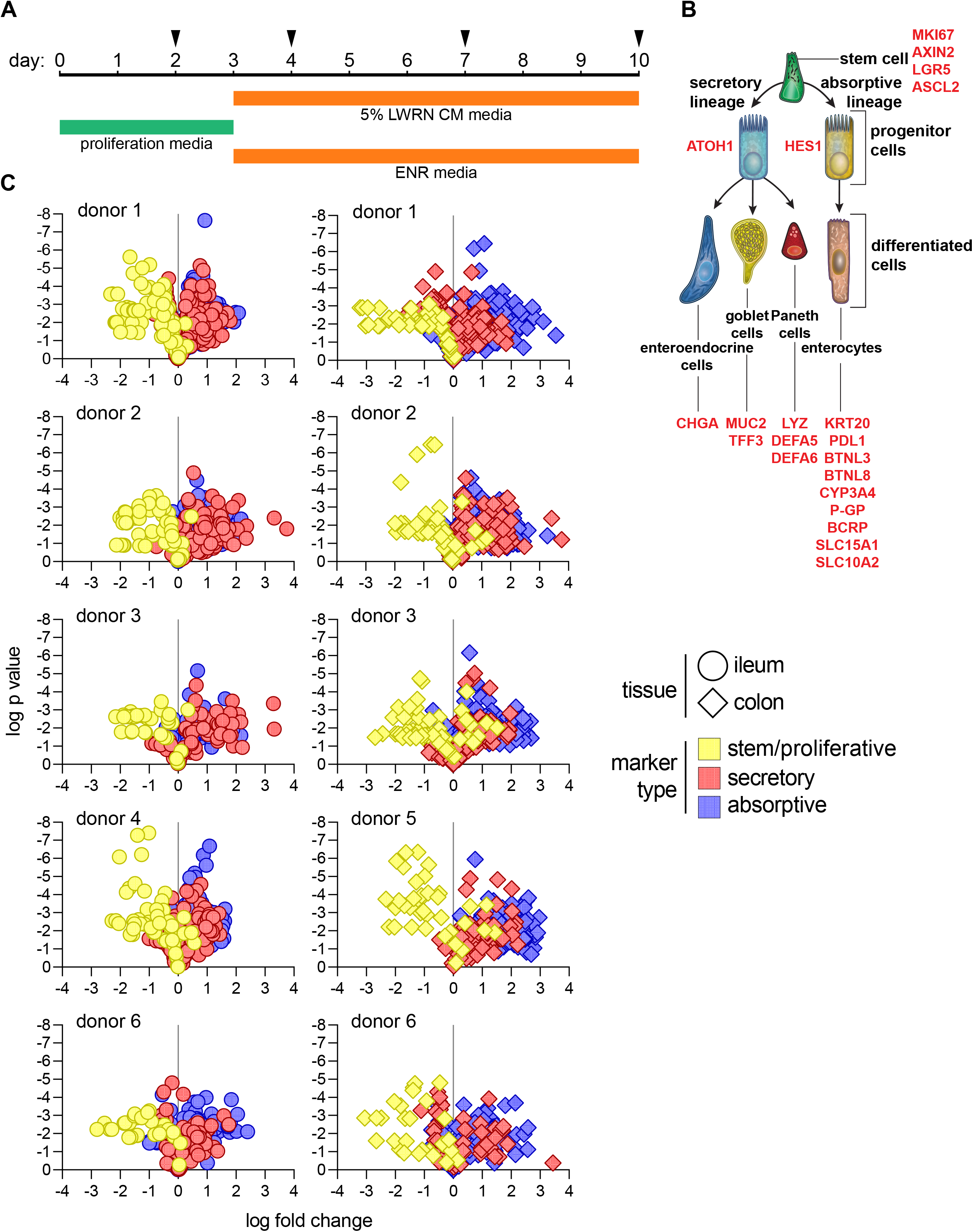
Consistent differentiation of ileum- and colon-derived ASC organoids (A) Schematic for time courses. Cultures were maintained for 10 days total, switching to differentiation media on day 3. Samples were collected on days 2, 4, 7, and 10. Day 2 cultures, maintained only in proliferation media, were used as a baseline for gene expression in subsequent analysis. (B) Stem/proliferative, secretory, and absorptive cell markers used to assess differentiation. (C) Assessing differentiation for each culture and donor. A ratio of gene expression (qPCR analysis) on day 4, 7, or 10 to gene expression on day 2 was calculated for each marker. Data were visualized as volcano plots for each culture, representing p value versus fold change in gene expression. Both ileum and colon cultures were present for all donors except for donor 4 (ileum only) and donor 5 (colon only). Each individual culture was analyzed using 2-3 independent time courses (details outlined in methods section); each time course included 3 cell culture (technical) replicates. Gray vertical lines represent fold change of 1. Circle symbols represent ileum cultures and diamond symbols represent colon cultures. Each symbol represents a marker for one sample (distinct replicate and time point). Colors represent each class of cell types (stem/proliferative, secretory, or absorptive) assessed.

Differentiation status of cultures was assessed using quantitative RT-PCR (qPCR), focusing on expression of developmental genes (Figure 1B). We designed a qPCR panel containing markers of stem/proliferative, secretory, and absorptive cells (Figure 1B). MKI67, LGR5, ASCL2, and AXIN2 were used as markers for proliferative and stem cells (Barker et al., 2007; Bullwinkel et al., 2006; Jho et al., 2002; Li et al., 2016; van der Flier et al., 2009). ATOH1 and HES1 were included as markers of secretory and absorptive progenitors, respectively (Gehart and Clevers, 2019). Additionally, markers for predominant functional secretory cell types were included: CHGA for enteroendocrine cells (Engelstoft et al., 2015), MUC2 and TFF3 for goblet cells (Pelaseyed et al., 2014), and LYZ, DEFA5, and DEFA6 for Paneth cells (Gassler, 2017). Several enterocyte markers were analyzed: CYP3A4, PGP, BCRP, SLC15A1, and SLC10A2 (International Transporter et al., 2010; Kaminsky and Zhang, 2003) were included as metabolic and absorptive markers and KRT20, PDL1, BTNL3, and BTNL8 were included as structural and intercellular signaling molecules (Di Marco Barros et al., 2016; Moll et al., 1990).

### Consistent developmental pathway gene expression patterns among donors

To assess differentiation potential of cultures, later time points (more differentiated) were compared to an early time point (more stem/proliferative). Transcript levels of the markers were examined, comparing days 4, 7, and 10 to day 2 and visualized on a volcano plot. This analysis showed that stem/proliferative cell markers were downregulated over time whereas secretory and absorptive cell markers were upregulated (Figure 1C). Importantly, ileum- and colon-derived cultures behaved similarly. This developmental pattern was shared among all cultures analyzed, indicating that differentiation potential was similar for all cultures.

The proliferation marker MKI67 was down regulated as cultures differentiated. Additionally, previously-described adult intestinal stem cell markers LGR5, ASCL2, and AXIN2 followed a similar pattern (Figure 2A). This pattern points to a shift to a mitotically arrested state over time, which is indicative of enrichment for terminally differentiated cells. ASCL2 gene expression was challenging to detect in all samples. Starting transcript levels were relatively low (day 2 C_t_ value 32.0 ± 2.46 for ASCL2 vs 25.9 ± 1.52 for LGR5, 23.6 ± 1.50 for AXIN2, and 21.8 ± 1.63 for MKI67 and Figure S2A), making reliable detection difficult as gene expression was downregulated over time. Therefore, this marker was excluded from subsequent analysis. Overall expression pattern for stem and proliferative cell markers was similar for ileum and colon cultures (Figure S2A). Comparing ASC organoids and original tissue, expression of stem and proliferative markers was similar (Figure S2A and S2D), although LGR5 expression was lower in ASC organoid cultures (average values in ileum: 0.0197 in ASC organoids, 0.302 in tissues; average values in colon: 0.0543 in ASC organoids, 0.167 in tissues).

**Figure 2:**
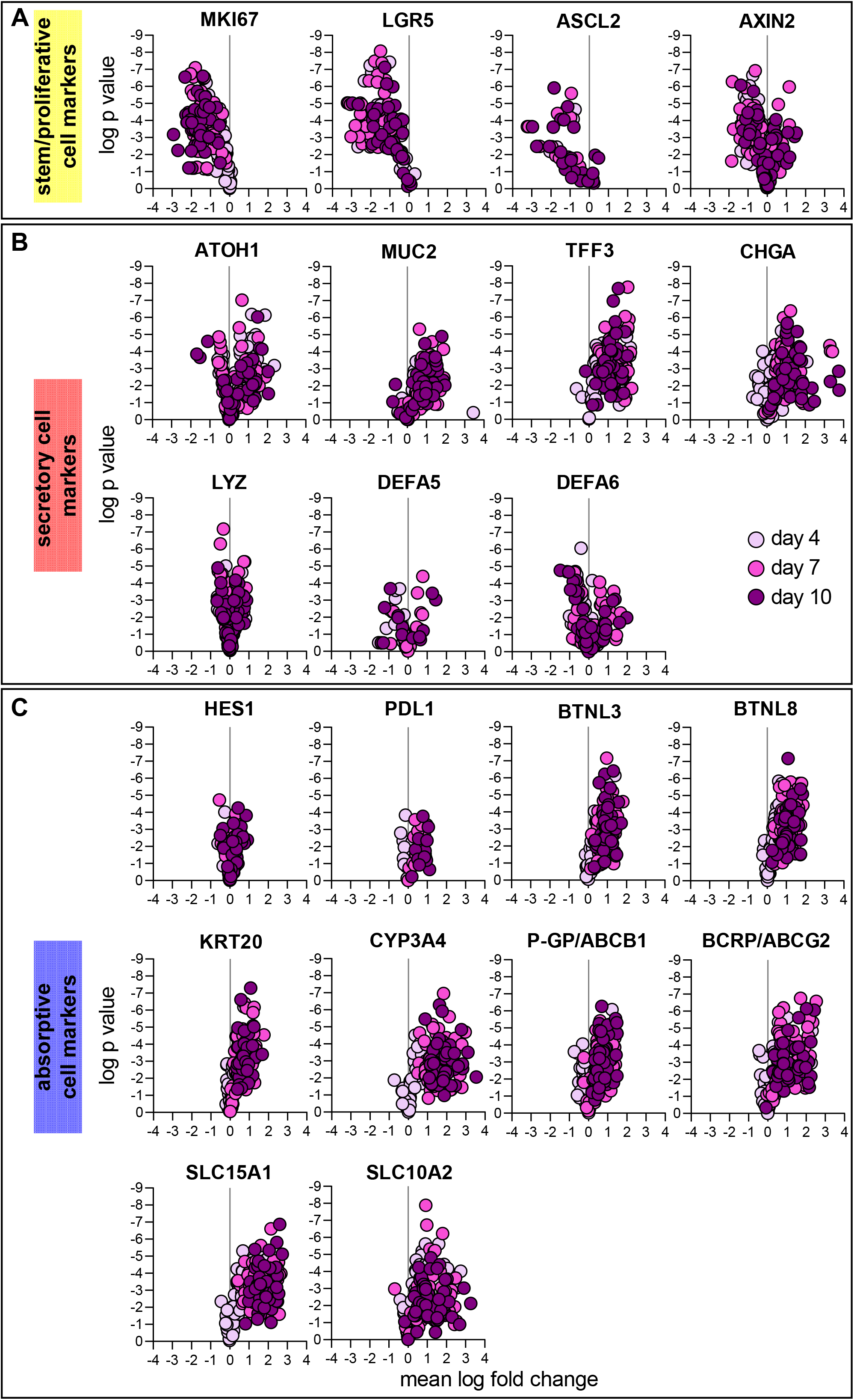
Regulation of stem/proliferative, secretory, and absorptive cell markers Expression of each stem/proliferative (A), secretory (B), and absorptive (C) marker is shown as a ratio of day 4, 7, or 10 to day 2 as was presented in Figure 1C. Plots are colored according to time point (darker colors are later time points) and each symbol represents a separate sample (distinct donor, replicate). As in Figure 1C, 2-3 time courses are presented for each culture and each time course experiment includes 3 cell culture (technical) replicates. All gene expression analysis was performed by qPCR.

Goblet and enteroendocrine cell markers MUC2, TFF3 and CHGA increased in expression as cultures developed, indicating the presence of these cells in well-differentiated cultures (Figure 2B). Expression of goblet cell markers was similar when comparing ileum and colon cultures (Figure S2B). Additionally, expression pattern in ASC organoids and human tissue was similar (Figure S2B and S2E). Increase in expression for Paneth cell markers LYZ, DEFA5, and DEFA6 was not dramatic. Appropriately, Paneth cell markers were detected at a lower level in colon cultures than ileum cultures (Figure S2B), consistent with the presence of Paneth cells in the small intestine but not colon (Clevers and Bevins, 2013). Tissue expression of Paneth cell markers was as expected: higher expression of all markers was observed in ileum (Figure S2E). Importantly, expression of DEFA5 and DEFA6 was higher in tissue than in ASC organoids. This result suggests that media conditions used were not conducive to the robust formation of Paneth cells, similar to previous results (Mead et al., 2018).

To assess the development of enterocytes, expression of several markers was determined. Structural and signaling molecules KRT20, PDL1, BTNL3, and BTNL8 increased expression over time (Figure 2C) and did not display distinct ileum-colon expression patterns (Figure S2C). Additionally, the expression of molecules important for metabolism and transport was quantified. Expression of CYP3A4 was examined as cytochrome P450 enzymes are important for intestinal metabolism. Expression of CYP3A4 increased dramatically over time in all cultures (Figure 2C). CYP3A4 expression was higher in ileum cultures than colon cultures (Figure S2C), reflecting the metabolic function of the small intestine, as has been observed previously (McKinnon et al., 1995). Additionally, the expression of ATP-binding cassette (ABC) family transporters P-GP and BCRP was upregulated over time (Figure 2C), showing little ileum-colon expression difference (Figure S2C). Similarly, expression of solute transporters SLC15A1 and SLC10A2 were upregulated considerably over time (Figure 2C). Expression of both solute transporters was higher in the ileum cultures (Figure S2C), once again emphasizing the importance of small intestine as the major site of nutrient transport. Interestingly, while expression levels for enterocyte markers was broadly similar in ASC organoids (Figure S2C) and original tissue (Figure S2F), ileum-colon polarity was lost for some transcripts. This was most apparent in transporters where P-GP, BCRP, and SLC15A1 were more highly expressed in ileum in original tissue but not in ASC organoids whereas SLC10A2 maintained its pattern of higher expression in ileum-derived ASC organoids.

### Quantifying inter- and intra-donor variability during epithelial cell differentiation

We determined the extent of variability among cultures using principal component and correlation analysis. Gene expression data generated by qPCR for developmental pathway analysis for both ASC organoids and original human tissue (discussed in Figures 1 and 2) was used. All data points were analyzed based on six features: tissue (ileum vs colon), donors, day of RNA harvest, media type, culture passage number, and replicates. Principal component analysis (PCA) showed that samples collected on each day of time course experiments clustered together (Figure 3A and 3B). Importantly, culture development and differentiation could be traced along principal component 1 (PC1), which captures most variation in data. Day 2 samples (highly proliferative/homogenous stem cultures) clustered away from day 7 and day 10 (highly differentiated/heterogeneous cultures) samples. Day 4 samples, which represent an intermediate state between proliferative/stem and differentiated cultures, fell between day 2 and day 7/10 samples along PC1. Additionally, day 7 and day 10 samples clustered closely, indicating that little further differentiation occurs after day 7. This analysis clearly showed that differentiated cultures were transcriptionally distinct from proliferative cultures. Importantly, original tissue samples clustered with day 2 ASC organoid samples (Figure 3A and 3B). Original tissue samples consisted of isolated crypts, where proliferative cells are located in the intestinal gland, making this observation biologically appropriate.

**Figure 3:**
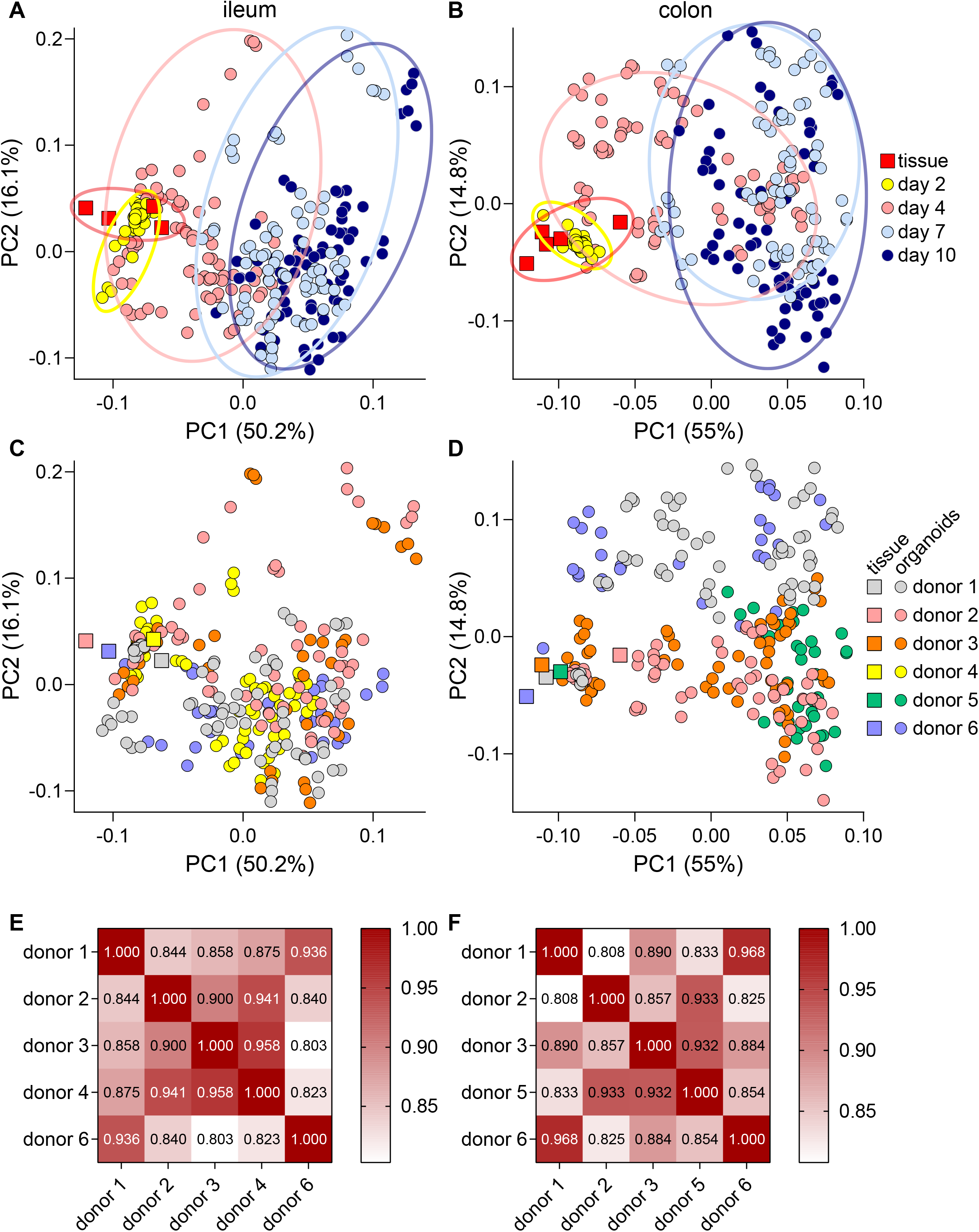
Quantifying variability among donors Principal component analysis of both ileum and colon ASC organoid cultures (circle symbols) and isolated human crypts (square symbols). Samples are marked by day of harvest (A and B) or donor identity (C and D). Percent variance for each principal component is reported in parentheses. Each symbol represents a separate RNA sample analyzed by qPCR. Correlation matrix (E and F) comparing gene expression patterns in differentiated cultures (day 7).

Variability among donors was examined next. On PCA plots, little clustering was observed when ileum and colon samples were colored based on originating donor (Figure 3C and 3D), suggesting that differences among donors is not a major driver of variability. Next, we sought to quantify the extent of similarity among ASC organoid cultures. To quantify similarity among donors, we employed Pearson correlation analysis. When comparing gene expression patterns in differentiated (day 7) ASC organoids, a high degree of correlation was observed (Figure 3E and 3F). Correlation coefficients ranged from 0.803 to 0.958 for ileum cultures and 0.808 to 0.968 for colon cultures. The high degree of correlation suggests that cultures broadly behave similarly when differentiated. This is an important outcome, indicating the likely generalizability of results produced in this culture system.

Additional variables among cultures were examined using PCA. As expected, samples incubated in the same media clustered together (Figure S3A). That is, samples grown in proliferation media clustered together and samples grown in either differentiation media clustered together. These results recapitulate findings from time-based PCA plots (Figure 3A and 3B). Additionally, this analysis shows that there is little difference between 5% LWRN CM and ENR differentiation media with respect to developmental pathways. Importantly, little clustering was observed when comparing samples over several passages and cell culture replicates suggesting minimal intra-donor variability (Figure S3B and S3C).

### Expression of inflammatory immune receptors is developmentally controlled

To evaluate the potential for modeling inflammatory diseases in ASC organoid cultures, the expression of immune receptors that initiate pro-inflammatory signaling pathways was quantified in proliferative and differentiated ASC organoid cultures. We focused on receptors for interferons (IFN) and tumor necrosis factor (TNF), as they have been implicated as important drivers of inflammatory gut indications (Andreou et al., 2020; Delgado and Brunner, 2019; Neurath, 2019). The expression of type I (IFNAR1 and IFNAR2), type II (IFNGR1 and IFNGR2), and type III (IFNLR1 and IL10R2) interferon receptors as well as two TNF receptors (TNFR1 and TNFR2) was quantified. Interestingly, receptor expression was higher in differentiated cultures compared with proliferative cultures (Figure 4A). This pattern was consistent among the three donors analyzed, suggesting that this phenomenon is generalizable. This pattern held true for type I (Figure 4B), type II (Figure 4C), and type III (Figure 4D) IFN receptors as well as TNF receptors (Figure 4E). These findings highlight the importance for thorough characterization of immune receptor expression prior to challenging cells with cytokines or pathogens.

**Figure 4:**
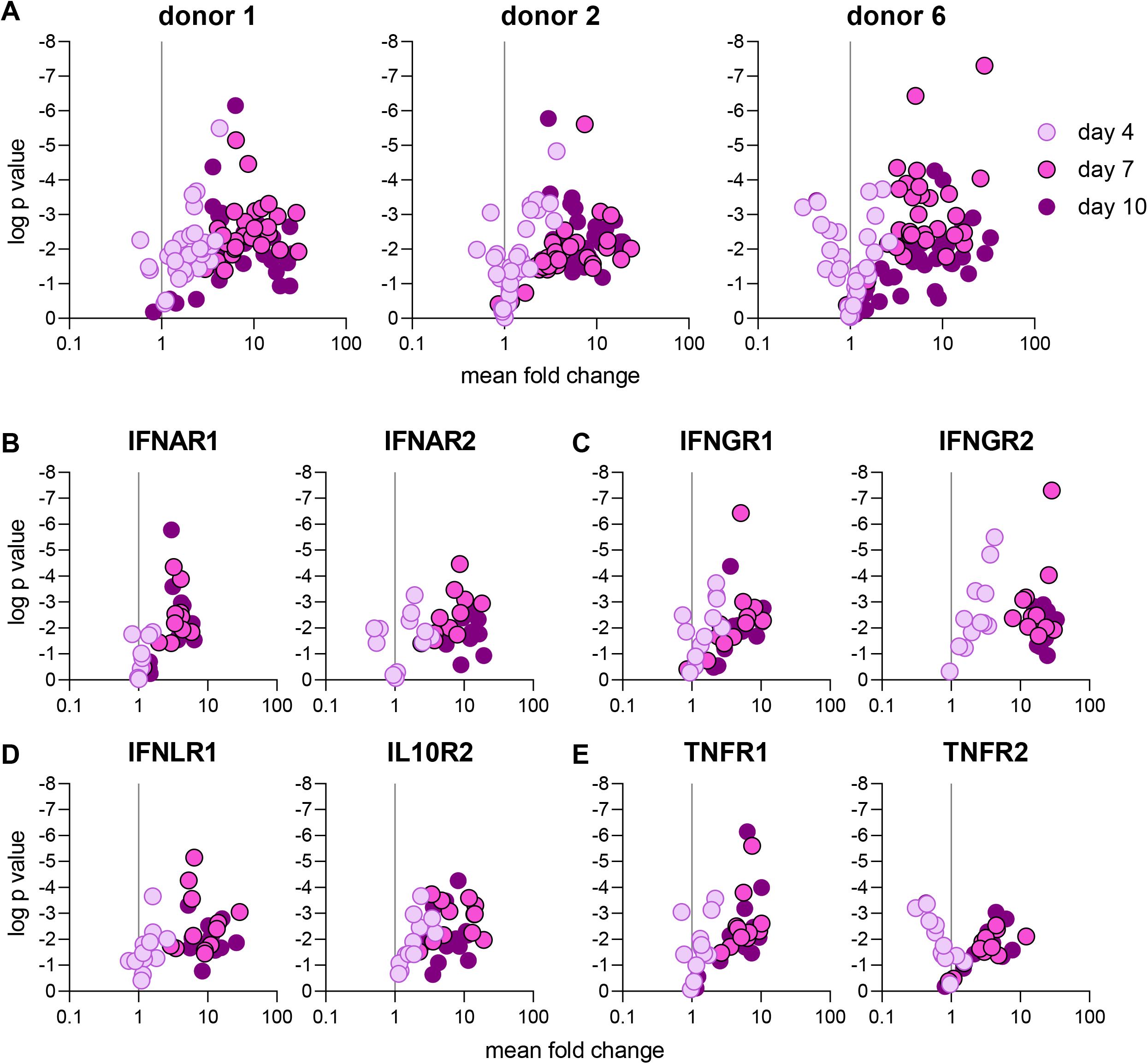
Expression of inflammatory immune receptors is developmentally controlled (A) Quantitative PCR was used to assess expression of IFN and TNF receptors in three donors and was displayed as a ratio of day 4, 7, or 10 to day 2 as was presented in Figure 1C. Plots are colored according to time point (darker colors are later time points) and each symbol represents a separate sample (distinct tissue, time point, and replicate). One time course per donor was included. Each time course included 3 cell culture (technical) replicates. (B-E) Expression of type I (B), type II (C), and type III (D) interferon and TNF (E) receptors, plotted in similar manner to Figure 2. Plots are colored according to time point (darker colors are later time points) and each symbol represents a separate sample (distinct donor, replicate).

IFN receptors did not show a statistically significant ileum-colon culture expression bias (Figure S4A-S4C). TNF receptors appeared to be more highly expressed in ileum cultures (Figure S4D). TNFR1 expression was indeed significantly higher in ileum cultures (p = 0.0008) whereas the difference for TNFR2 expression was not statistically significant (p = 0.0997). Robust and reproducible expression of IFN and TNF receptors indicate that this ASC organoid system is suitable for modeling intestinal inflammatory disease *in vitro*.

### Markers of functional intestinal cells are detected at protein level

The presence and fractions of functional cell types in ASC organoid cultures were determined using immunofluorescence microscopy (Figure 5). Differentiated cultures (day 7) were fixed and markers of Paneth (lysozyme), goblet (mucin 2), and enteroendocrine (chromogranin A) cells were detected. Structural proteins actin and E-cadherin were detected as markers of apical microvilli and cell junctions, respectively. This analysis showed that ASC organoids formed polarized structures with apical surfaces facing the lumen, as expected. Additionally, uniform E-cadherin staining was detected, indicating the formation of tight barrier surfaces. Finally, Lysozyme, mucin 2, and chromogranin A were detected in cultures, confirming the presence of secretory cells in day 7 cultures (Figure 5A). Importantly, brightfield imaging confirmed the presence of a healthy epithelium as cultures differentiated (Figure S5). Next, the fraction of each cell type was quantified in both ileum- and colon-derived differentiated ASC organoids from several (n = 3) donors (Figure 5B). Goblet cell marker mucin 2 and enteroendocrine cell marker chromogranin A were detected at relatively high rates whereas Paneth cell marker lysozyme occurred at much lower rates. These data indicate that while culture conditions are conducive to the formation of goblet and enteroendocrine cells, additional pathway modulators would have to be included in the culture media to increase the proportions of Paneth cells. Importantly, the staining patterns and cell proportions were consistent among the three donors examined indicating that differentiation patterns among ASC organoid cultures are uniform.

**Figure 5:**
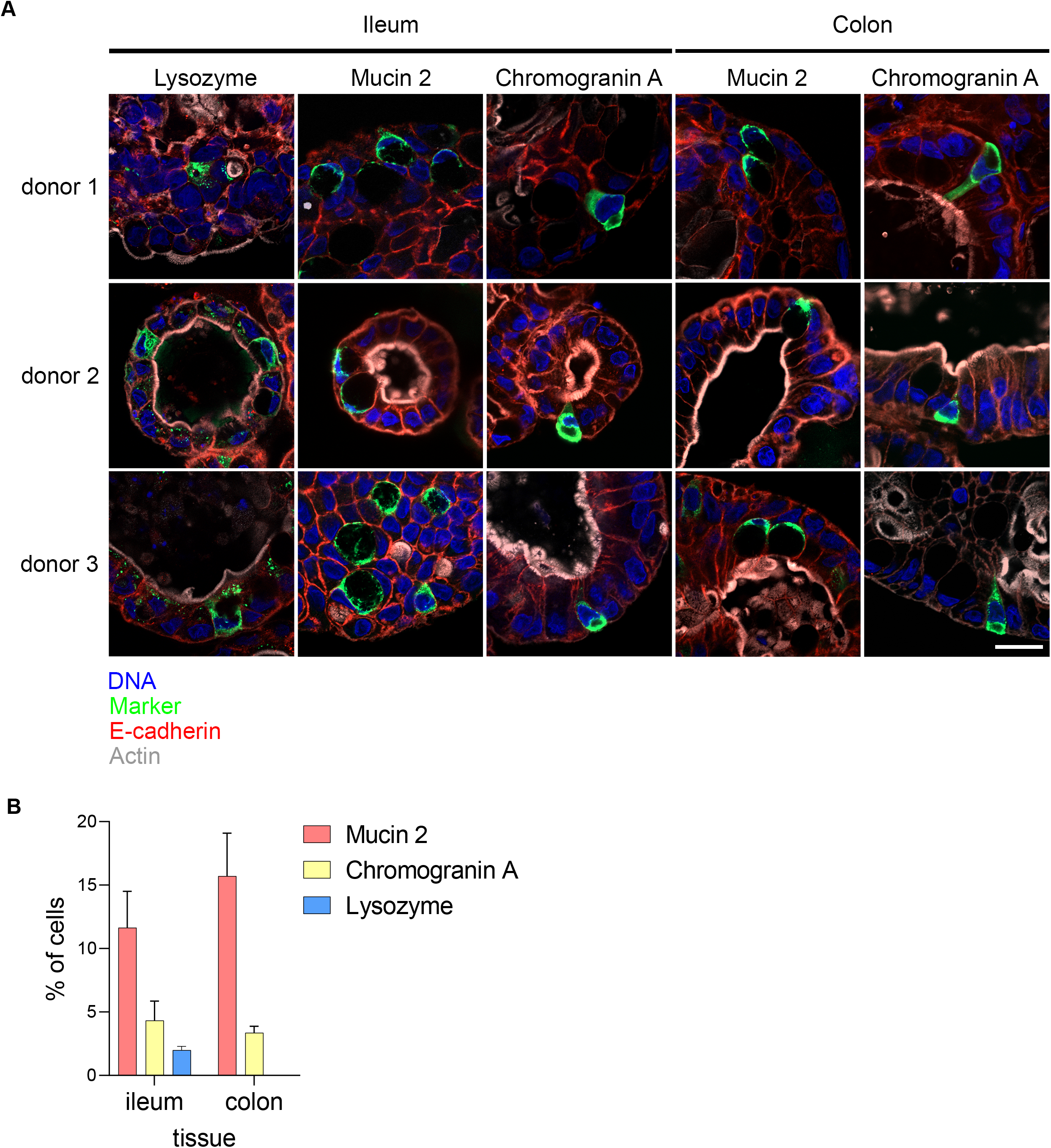
Immunofluorescence imaging to confirm presence of differentiated cells (A) Differentiated ASC organoids (harvested at day 7 as outlined in Figure 1) from three donors were fixed and markers of mature intestinal cells were detected using confocal microscopy. Pseudocoloring is as follows: Blue: DNA; Green: lysozyme, mucin 2, or Chromogranin A; Red: E-cadherin; Gray: actin. Scale bar = 20μm (applies to all panels). (B) Quantification of cell type markers. Differentiated (day 7) Ileum and colon cultures from three donors were stained for lysozyme (Paneth cell marker, ileum cultures only), mucin 2 (goblet cell marker, ileum and colon cultures), chromogranin A (enteroendocrine cells, ileum and colon cultures), and DNA. Fraction of each cell type present was calculated by counting marker-positive cells among hundreds of total cells (>500 cells for all samples and markers except for lysozyme staining in donor 2 (307 cells) and donor 3 (473 cells) cultures). Data are presented as percent of total cells positive for a given cell type marker.

### Ileum- and colon-derived ASC organoids secrete serotonin, GLP1, and PYY

The intestine is an important endocrine tissue and gut hormones play key roles in satiety, glucose homeostasis, and nutrient absorption (Ahlman and Nilsson, 2001; Martin et al., 2019; Worthington et al., 2018). However, cell culture systems to study gut hormone production and secretion *in vitro* have not been readily available. Thus, we quantified hormone secretion by differentiated ileum and colon ASC organoids (day 7 post seeding, stimulated as described in methods section) derived from three donors. As enteroendocrine cell markers were robustly detected (Figure 2 and 5), we decided to assess function by measuring levels of secreted serotonin, PYY, and GLP1. The secretion of all three hormones was detected for all donors (Figure 6). Serotonin secretion was consistently detected more highly in colon cultures than ileum cultures (Figure 6A). Compared with untreated cultures, serotonin secretion increased by 1.1-fold (ileum) or 7.5-fold (colon). GLP1 secretion was robustly detected in all cultures as well (Figure 6B). While both ileum and colon cultures secreted GLP1, the magnitude of secretion varied. Ileum cultures displayed an average 7.4-fold induction over untreated cultures while colon cultures displayed a 22.7-fold increase. Interestingly, the basal levels of GLP1 were lower in colon cultures than ileum cultures. Finally, secretion of PYY was detected in all cultures (Figure 6C). While the basal and stimulated levels of PYY were higher in colon than ileum, the magnitude of induction upon stimulation was similar (5.7-fold for both ileum and colon cultures). This is consistent with higher colon PYY expression levels *in vivo* (Zhou et al., 2006). Importantly, all three donors analyzed in this experiment behaved similarly, suggesting that enteroendocrine cell function is consistently observed in ASC organoid cultures. Taken together these data demonstrate that ASC organoid cultures faithfully recapitulate physiological conditions for hormone secretion.

**Figure 6:**
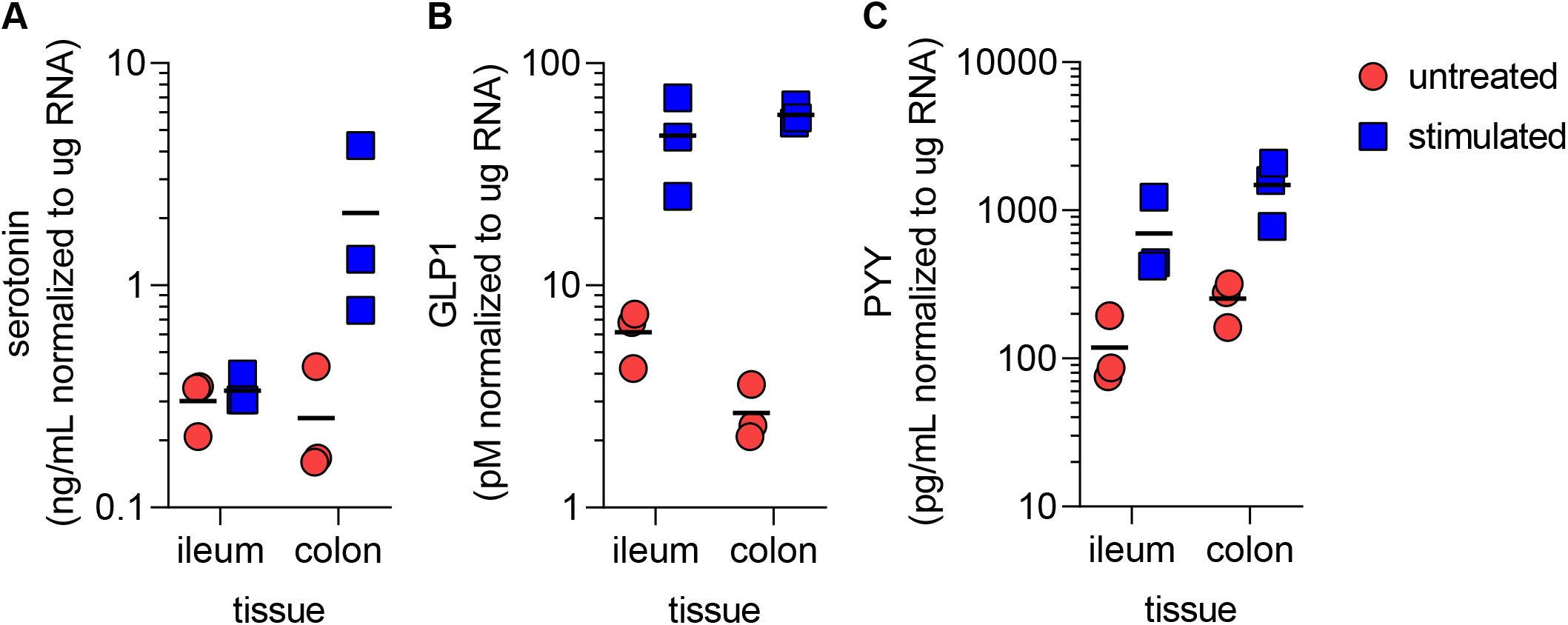
Ileum- and colon-derived ASC organoids secrete serotonin, GLP1, and PYY Secretion of serotonin (A), GLP1 (B), and PYY (C) by ileum- and colon-derived ASC organoid cultures. Cultures were stimulated overnight by 10μM forskolin, 10μM IBMX, and 10mM glucose and hormone amounts were quantified by ELISA. Cultures from 3 donors were analyzed and experiments were performed with 4 cell culture (technical) replicates. Each symbol in the figure represents the average of cell culture replicates for each donor culture analyzed.

### Central carbon metabolism changes during ASC organoid differentiation

Central carbon metabolism is a complex enzyme-mediated network that converts sugars into precursors for metabolism. Targeted metabolomics analysis of the ileum- and colon-derived ASC organoids assessed the metabolic differences of differentiated ASC organoids compared to stem cell-enriched cultures. Using triple quadrupole (QQQ) liquid chromatography/mass spectrometry (LC/MS), 108 central carbon metabolites were profiled at 3 timepoints (2, 4, and 7 days after initial plating) for ileum- and colon-derived ASC organoids from 5 and 4 donors, respectively. For each tissue type, metabolite concentrations were compared between donors at each timepoint. PCA results (Figure 7A and B) indicated minimal metabolic donor-to-donor variability as donors for a particular timepoint tended to cluster together. This robust clustering pattern indicated that the metabolic state of cultures is consistent. Clear differences in metabolism were evident between each day, as indicated by separated clusters for each timepoint. These metabolic trends were consistent in both ileum- and colon-derived ASC organoids.

**Figure 7:**
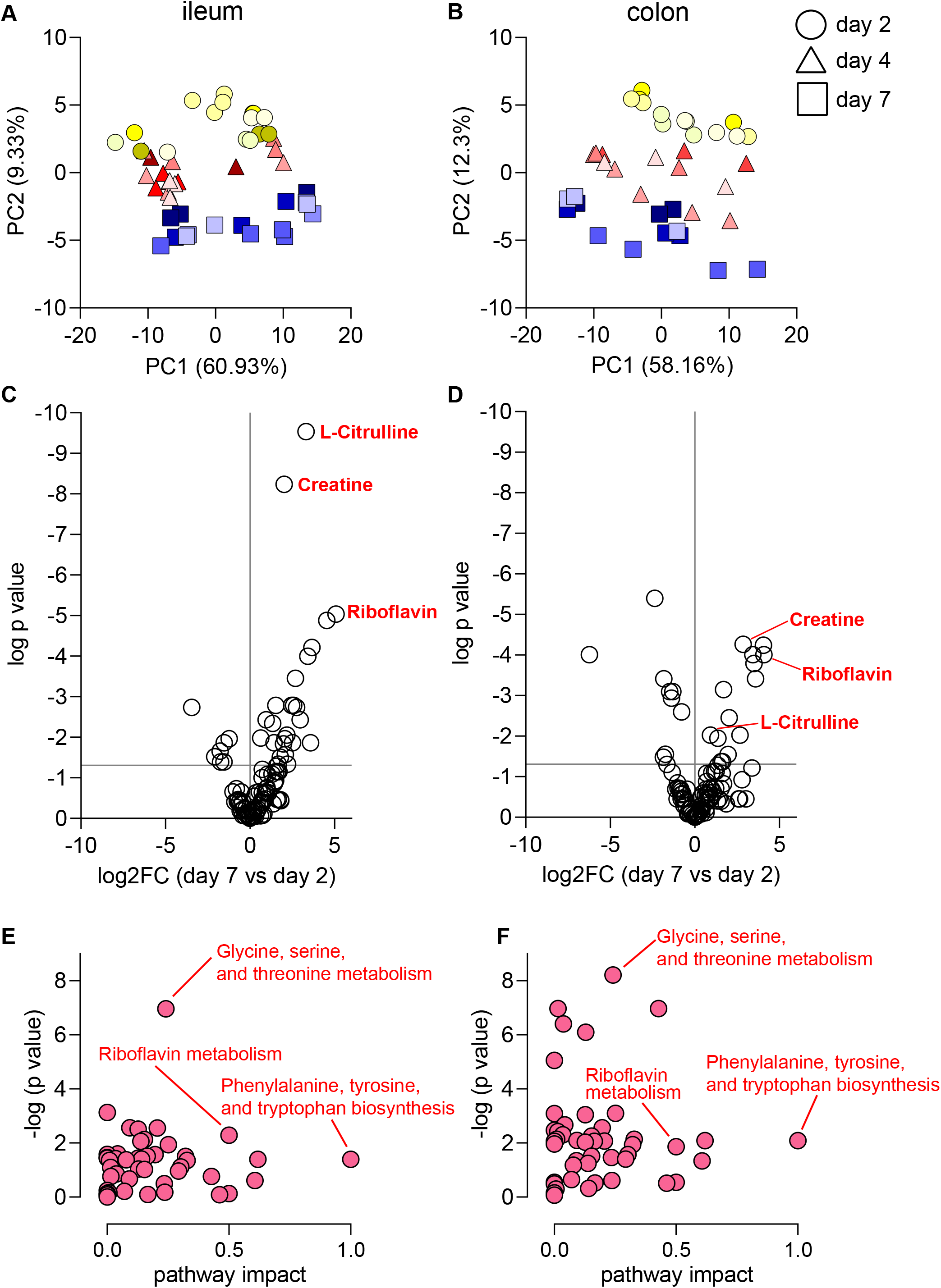
Metabolomics analysis of ASC organoid cultures (A and B) Principal component analysis of central carbon metabolites in ileum- and colon-derived ASC organoids. Percent variance for each principal component is reported in parentheses. Each symbol represents a cell culture replicate and shading represents different donors. Symbols with the same shading are cell culture (technical) replicates. 5 ileum culture donors and 4 colon culture donors are represented, each containing 3 cell culture (technical) replicates. (C and D) Abundance of metabolites presented as ratio of day 7 abundance to day 2 abundance for ileum- and colon-derived ASC organoids. (E and F) Pathway analysis for metabolite abundance with key pathways highlighted. Panels A, C, and E represent ileum cultures and panels B, D, and F represent colon cultures.

To better understand the changes in central carbon metabolism during ASC organoid development, significantly different metabolites (p < 0.05, one-way ANOVA) were visualized by volcano plot (Figure 7C and D). L-citrulline, creatine, riboflavin, and L-glutathione (oxidized) were among the list of significantly differential metabolites between Day 2 and 7. Enrichment analysis (Figure 7E and F) revealed that amino acid metabolism – specifically glycine/serine/threonine pathway and arginine/proline pathway – was affected by differentiation to mature ileum- and colon-derived ASC organoids.

The change in abundance of several of these metabolites is consistent with metabolism in these tissues in the human body. For example, the relative abundance of L-citrulline increased by nearly 10 times in ileum-derived ASC organoids from Day 2 to Day 7. This result is consistent with previous studies, which have demonstrated a role for citrulline in intestinal function, and that absorption of citrulline occurs in the middle to lower ileum (Fragkos and Forbes, 2018; Vadgama and Evered, 1992). Riboflavin, which significantly increased from Day 2 to Day 7 in both colon and ileum-derived ASC organoids, has previously been shown to be absorbed in the small intestine (Feder et al., 1991; Hegazy and Schwenk, 1983). Flavin adenine dinucleotide (FAD), which is in the same metabolic pathway as riboflavin, also increased in both ASC organoid types. Taken together, our data suggest that the metabolic profiles detected in these ASC organoid cultures are consistent with what is known about their physiologic functions. Nevertheless, to more precisely determine the ability of these ASC organoids to recapitulate tissue metabolism, additional experiments are warranted.

## Discussion

The advent of three-dimensional adult stem cell (ASC) organoid models allows for *in vitro* modeling of human tissue providing a more physiologically relevant system than traditional tissue culture. One concern about these systems is the level of variability that may be observed from culture to culture. Here we described the characterization of ileum and colon ASC organoids derived from various donors. We have demonstrated similar levels of epithelial cell differentiation amongst donors regardless of media used for differentiation, or passage number of the culture. Critically, donor-to-donor variability was minimal.

The ASC organoid cultures examined here differentiate on a time-scale that matches the human gut epithelium. Within a few days, differentiation is complete, as indicated by the overlap between gene expression signatures at day 7 and 10 (Figure 3). Importantly, only markers of enterocytes, goblet, Paneth, and enteroendocrine cells could be detected (Figures 1, 2, and 5); rarer cell types such as tuft and microfold (M) cell markers were not detected (data not shown). This is consistent with previous work showing that differentiation toward rare cell type linages requires the addition of specific cytokines (Boonekamp et al., 2020).

Curiously, expression of Paneth cell markers was observed in colon cultures (Figure S2). While Paneth cell marker expression levels were higher in ileum cultures, colon culture expression was not absent. This observation could be explained by the complex nature of marker expression. Recent single-cell analysis publications (Parikh et al., 2019; Smillie et al., 2019; Wang et al., 2020) have highlighted the fact that gradients of marker expression exist in each cell type. Additionally, Paneth-like cells have been identified in the human colon (Wang et al., 2020), indicating that Paneth cell marker expression may represent this cell type. Overall, Paneth cell differentiation was not very efficient in our organoid cultures (Figure 2, S2, and 5). It is known that both Wnt and Notch signaling pathways control Paneth cell differentiation (Sato et al., 2011b). The differentiation media conditions used in this study did not include Wnt and Notch modulators, which likely explains the modest increase in Paneth cell markers. While Paneth cells are observed in low quantities (Figure 5), increased representation of this cell type would be enhanced by adding exogenous Wnt agonists and Notch antagonists to the media as previously described (Mead et al., 2018; Yin et al., 2014).

We show that differentiation is remarkably consistent among cultures. Appropriately, gene expression patterns are fairly uniform at early time points (denoted by a tight cluster colored in yellow – Figure 3A and 3B), reflecting homogeneity of stem/proliferative-cells in the intestine. As cultures differentiate over time, gene expression patterns become less uniform (pink and blue symbols – Figure 3A and 3B), which is indicative of expected cell type heterogeneity in terminally differentiated intestinal cells. This pattern is reminiscent of the “epigenetic landscape” described by Waddington (Goldberg et al., 2007). Our data nicely depict the widening of cell fate trajectories as cultures differentiate to generate functional cells.

Our gene expression analysis highlighted region-specific expression patterns for certain transcripts. These included Paneth cell markers LYZ, DEFA5, and DEFA6 and enterocyte markers CYP3A4, P-GP, BCRP, SLC15A1, and SLC10A2 (Figure S2E-F). This pattern represents expected intestinal biology: Paneth cells are not typically present in the colon (Clevers and Bevins, 2013; Gassler, 2017) and the small intestine is the major site for transport and first pass metabolism (International Transporter et al., 2010; Kato, 2008). It has been shown previously that regional gene expression patterns are maintained in gut organoid cultures (Middendorp et al., 2014). We observed that regional gene expression patterns were largely maintained for transcripts that displayed an ileum-colon expression bias. However, some transcripts lost ileum-colon expression bias (transporters P-GP, BCRP, and SLC15A1). These results suggest that regional expression patterns *in vivo* are governed by specific factors that are absent in a neutral selection condition such as media used to culture ASC organoids. Factors controlling regional expression patterns have not been defined but could include signals from the luminal microbial community, immune cells in lamina propria, or other to-be-defined signals originating from serosal sources.

Large variability in primary culture systems could lead to uninterpretable results. This requires meticulous quantification of various sources of variability, leading to an understanding of noise in the experimental system. We found that among several variables – donors, differentiation media used, passages, and cell culture replicates – variability was manageable (Figure 3). This information indicates that cultures behave similarly during the differentiation process and over multiple passages, thereby enabling robust experimental design.

Hormone production and secretion is a critical function of the human intestine, so we investigated endocrine cell function in ASC organoids. The gut is often referred to as the largest endocrine tissue in the human body (Ahlman and Nilsson, 2001), producing many hormones that regulate a range of metabolic functions in the intestine and throughout the body. The ASC organoid cultures examined for this work effectively produced and secreted GLP-1 and PYY (Figure 6), which work in concert with pancreatic hormones to regulate blood glucose levels (Gribble and Reimann, 2019; Worthington et al., 2018). Additionally, we show that ASC organoid cultures produced and secreted serotonin (Figure 6). Serotonin plays important roles in the intestine by regulating peristalsis, vasodilation, and pain perception (Mawe and Hoffman, 2013; Spohn and Mawe, 2017). In addition to regulating metabolic and intestinal functions, these hormones have been implicated in various diseases. For example, GLP-1, PYY, and serotonin levels change during inflammatory bowel disease (IBD) and serotonin is thought to act as a proinflammatory molecule in celiac disease (Worthington et al., 2018). Furthermore, recent work has uncovered a novel mechanism through which intestinal microbes induce serotonin secretion (Sugisawa et al., 2020). Intestinal ASC organoids provide a robust *ex vivo* platform to identify modulators of endocrine cell function. These modulators will be important for studying disease processes as well as basic intestinal function.

We observed differential production of central carbon metabolites at various time points of culture differentiation. As the cultures differentiated, there was an increase in L-citrulline, creatine and riboflavin. This represents the metabolites being utilized by the cells present in this heterogenous population. While this targeted approach yields interesting reproducible information about central carbon metabolism, future studies could take an untargeted approach to see if additional metabolites can be detected in a reproducible manner. The results could indicate the ratio of a particular cell population or unique metabolic pathways that play important roles in the intestine.

In this work we have characterized variability in primary human intestinal ASC organoids and demonstrated that donor-to-donor variation is minimal. Additionally, we have shown that the cell culture system recapitulates key features of the originating tissue. Identifying and characterizing the cell state for primary cell cultures is important for developing robust applications. Without thorough characterization, assay results will be difficult to interpret. Put together, thorough characterization of cell state and functional cell information lead to successful application of this primary cell platform to ask fundamental biological questions as well as to advance therapeutic discovery.

## Experimental Procedures

For complete Experimental Procedures see supplemental materials.

### ASC organoid culture growth and differentiation

Intestinal organoid cultures were established as previously described (Miyoshi and Stappenbeck, 2013; Sato et al., 2011a). Cultures were maintained in proliferation media (50% LWRN conditioned media + Y27632 + SB431542) for routine passaging. To induce differentiation, culture media was changed to either a serum-containing media (5% LWRN conditioned media, no Y27632 or SB431542) or a defined media (base medium containing EGF, Noggin, and R-spondin) three days after organoids were plated.

### Transcriptional and immunofluorescence microscopy analysis

At indicated time points, cells were harvested for RNA purification and quantitative PCR analysis. For immunofluorescence analysis, organoids were removed from Matrigel using cell recovery solution (Corning) and fixed using 4% paraformaldehyde. After permeabilization and blocking, organoids were incubated overnight in antibodies targeting differentiated cell markers, followed by secondary antibody staining. Stained organoids were mounted onto slides and imaged using confocal microscopy.

### Principal component and correlation analysis

Principal component analysis was carried out using R. The results were visualized using Graphpad Prism. Pearson correlation analysis was performed using R, including only samples from the day 7 time point. The correlation matrix was visualized using Graphpad Prism.

### Hormone secretion assay

Hormone secretion was induced in differentiated organoids (7 days post plating). Briefly, secretion was induced by adding forskolin, IBMX, and glucose to culture media and analyzing supernatants after overnight incubation using kits to detect secreted PYY, GLP1, and serotonin.

### Metabolomics analysis

Metabolites in organoids cultured for 2, 4, or 7 days were analyzed by mass spectrometry. Organoids were removed from Matrigel using Cell Recovery Solution (Corning) and extracted with acetonitrile/methanol/water. After homogenization, organoids were extracted for 10 minutes at −20°C and centrifuged. Supernatants were dried under nitrogen then dissolved in water/methanol for mass spectrometry analysis.

## Acknowledgments

We thank Erik Hett, Alex Therien, and Daria Hazuda for constructive comments on manuscript and continued support. We thank Sharon O’Brien for illustration in Figure 1B.

## Author contributions

Conceptualization, SM and LAL; Methodology, SM, TPW and CHW; Software and Formal Analysis, CHW; Investigation, SM, CMP, CWW, TPW, LAL; Data Curation, SM, TPW, CHW, TRS, LAL; Writing – Original Draft, SM and LAL; Writing – Review & Editing, all authors; Visualization, SM, TPW, CHW, TRS; Supervision, LAL; Project Administration, SM and LAL.

## References

Ahlman, H., and Nilsson (2001). The gut as the largest endocrine organ in the body. Ann Oncol 12 Suppl 2, S63–68.

Andreou, N.P., Legaki, E., and Gazouli, M. (2020). Inflammatory bowel disease pathobiology: the role of the interferon signature. Ann Gastroenterol 33, 125–133.

Barker, N., van Es, J.H., Kuipers, J., Kujala, P., van den Born, M., Cozijnsen, M., Haegebarth, A., Korving, J., Begthel, H., Peters, P.J., et al. (2007). Identification of stem cells in small intestine and colon by marker gene Lgr5. Nature 449, 1003–1007.

Bartfeld, S. (2016). Modeling infectious diseases and host-microbe interactions in gastrointestinal organoids. Dev Biol 420, 262–270.

Boonekamp, K.E., Dayton, T.L., and Clevers, H. (2020). Intestinal organoids as tools for enriching and studying specific and rare cell types: advances and future directions. J Mol Cell Biol.

Bullwinkel, J., Baron-Luhr, B., Ludemann, A., Wohlenberg, C., Gerdes, J., and Scholzen, T. (2006). Ki-67 protein is associated with ribosomal RNA transcription in quiescent and proliferating cells. J Cell Physiol 206, 624–635.

Chen, C., Yin, Y., Tu, Q., and Yang, H. (2018). Glucose and amino acid in enterocyte: absorption, metabolism and maturation. Front Biosci (Landmark Ed) 23, 1721–1739.

Clevers, H.C., and Bevins, C.L. (2013). Paneth cells: maestros of the small intestinal crypts. Annu Rev Physiol 75, 289–311.

Date, S., and Sato, T. (2015). Mini-gut organoids: reconstitution of the stem cell niche. Annu Rev Cell Dev Biol 31, 269–289.

Dawson, P.A. (2011). Role of the intestinal bile acid transporters in bile acid and drug disposition. Handb Exp Pharmacol, 169–203.

Delgado, M.E., and Brunner, T. (2019). The many faces of tumor necrosis factor signaling in the intestinal epithelium. Genes Immun 20, 609–626.

Di Marco Barros, R., Roberts, N.A., Dart, R.J., Vantourout, P., Jandke, A., Nussbaumer, O., Deban, L., Cipolat, S., Hart, R., Iannitto, M.L., et al. (2016). Epithelia Use Butyrophilin-like Molecules to Shape Organ-Specific gammadelta T Cell Compartments. Cell 167, 203–218 e217.

Dillon, A., and Lo, D.D. (2019). M Cells: Intelligent Engineering of Mucosal Immune Surveillance. Front Immunol 10, 1499.

Dubey, R., van Kerkhof, P., Jordens, I., Malinauskas, T., Pusapati, G.V., McKenna, J.K., Li, D., Carette, J.E., Ho, M., Siebold, C., et al. (2020). R-spondins engage heparan sulfate proteoglycans to potentiate WNT signaling. Elife 9.

Engelstoft, M.S., Lund, M.L., Grunddal, K.V., Egerod, K.L., Osborne-Lawrence, S., Poulsen, S.S., Zigman, J.M., and Schwartz, T.W. (2015). Research Resource: A Chromogranin A Reporter for Serotonin and Histamine Secreting Enteroendocrine Cells. Mol Endocrinol 29, 1658–1671.

Feder, S., Daniel, H., and Rehner, G. (1991). In vivo kinetics of intestinal absorption of riboflavin in rats. J Nutr 121, 72–79.

Fish, E.M., and Burns, B. (2020). Physiology, Small Bowel. In StatPearls (Treasure Island (FL)).

Fragkos, K.C., and Forbes, A. (2018). Citrulline as a marker of intestinal function and absorption in clinical settings: A systematic review and meta-analysis. United European Gastroenterol J 6, 181–191.

Gao, G., Li, J., Zhang, Y., and Chang, Y.Z. (2019). Cellular Iron Metabolism and Regulation. Adv Exp Med Biol 1173, 21–32.

Gassler, N. (2017). Paneth cells in intestinal physiology and pathophysiology. World J Gastrointest Pathophysiol 8, 150–160.

Gehart, H., and Clevers, H. (2019). Tales from the crypt: new insights into intestinal stem cells. Nat Rev Gastroenterol Hepatol 16, 19–34.

Goldberg, A.D., Allis, C.D., and Bernstein, E. (2007). Epigenetics: a landscape takes shape. Cell 128, 635–638.

Gribble, F.M., and Reimann, F. (2019). Function and mechanisms of enteroendocrine cells and gut hormones in metabolism. Nat Rev Endocrinol 15, 226–237.

Hegazy, E., and Schwenk, M. (1983). Riboflavin uptake by isolated enterocytes of guinea pigs. J Nutr 113, 1702–1707.

Hernando, N., and Wagner, C.A. (2018). Mechanisms and Regulation of Intestinal Phosphate Absorption. Compr Physiol 8, 1065–1090.

Hussain, M.M. (2014). Intestinal lipid absorption and lipoprotein formation. Curr Opin Lipidol 25, 200–206.

International Transporter, C., Giacomini, K.M., Huang, S.M., Tweedie, D.J., Benet, L.Z., Brouwer, K.L., Chu, X., Dahlin, A., Evers, R., Fischer, V., et al. (2010). Membrane transporters in drug development. Nat Rev Drug Discov 9, 215–236.

Jho, E.H., Zhang, T., Domon, C., Joo, C.K., Freund, J.N., and Costantini, F. (2002). Wnt/beta-catenin/Tcf signaling induces the transcription of Axin2, a negative regulator of the signaling pathway. Mol Cell Biol 22, 1172–1183.

Johansson, M.E., Sjovall, H., and Hansson, G.C. (2013). The gastrointestinal mucus system in health and disease. Nat Rev Gastroenterol Hepatol 10, 352–361.

Kaminsky, L.S., and Zhang, Q.Y. (2003). The small intestine as a xenobiotic-metabolizing organ. Drug Metab Dispos 31, 1520–1525.

Kato, M. (2008). Intestinal first-pass metabolism of CYP3A4 substrates. Drug Metab Pharmacokinet 23, 87–94.

Ko, C.W., Qu, J., Black, D.D., and Tso, P. (2020). Regulation of intestinal lipid metabolism: current concepts and relevance to disease. Nat Rev Gastroenterol Hepatol 17, 169–183.

Li, N., Yousefi, M., Nakauka-Ddamba, A., Tobias, J.W., Jensen, S.T., Morrisey, E.E., and Lengner, C.J. (2016). Heterogeneity in readouts of canonical wnt pathway activity within intestinal crypts. Dev Dyn 245, 822–833.

Lindemans, C.A., Calafiore, M., Mertelsmann, A.M., O’Connor, M.H., Dudakov, J.A., Jenq, R.R., Velardi, E., Young, L.F., Smith, O.M., Lawrence, G., et al. (2015). Interleukin-22 promotes intestinal-stem-cell-mediated epithelial regeneration. Nature 528, 560–564.

Martin, A.M., Sun, E.W., and Keating, D.J. (2019). Mechanisms controlling hormone secretion in human gut and its relevance to metabolism. J Endocrinol 244, R1–R15.

Mawe, G.M., and Hoffman, J.M. (2013). Serotonin signalling in the gut--functions, dysfunctions and therapeutic targets. Nat Rev Gastroenterol Hepatol 10, 473–486.

McKinnon, R.A., Burgess, W.M., Hall, P.M., Roberts-Thomson, S.J., Gonzalez, F.J., and McManus, M.E. (1995). Characterisation of CYP3A gene subfamily expression in human gastrointestinal tissues. Gut 36, 259–267.

Mead, B.E., Ordovas-Montanes, J., Braun, A.P., Levy, L.E., Bhargava, P., Szucs, M.J., Ammendolia, D.A., MacMullan, M.A., Yin, X., Hughes, T.K., et al. (2018). Harnessing single-cell genomics to improve the physiological fidelity of organoid-derived cell types. BMC Biol 16, 62.

Michels, B.E., Mosa, M.H., Streibl, B.I., Zhan, T., Menche, C., Abou-El-Ardat, K., Darvishi, T., Czlonka, E., Wagner, S., Winter, J., et al. (2020). Pooled In Vitro and In Vivo CRISPR-Cas9 Screening Identifies Tumor Suppressors in Human Colon Organoids. Cell Stem Cell 26, 782–792 e787.

Middendorp, S., Schneeberger, K., Wiegerinck, C.L., Mokry, M., Akkerman, R.D., van Wijngaarden, S., Clevers, H., and Nieuwenhuis, E.E. (2014). Adult stem cells in the small intestine are intrinsically programmed with their location-specific function. Stem Cells 32, 1083–1091.

Miyoshi, H., and Stappenbeck, T.S. (2013). In vitro expansion and genetic modification of gastrointestinal stem cells in spheroid culture. Nat Protoc 8, 2471–2482.

Moll, R., Schiller, D.L., and Franke, W.W. (1990). Identification of protein IT of the intestinal cytoskeleton as a novel type I cytokeratin with unusual properties and expression patterns. J Cell Biol 111, 567–580.

Neurath, M.F. (2019). Targeting immune cell circuits and trafficking in inflammatory bowel disease. Nat Immunol 20, 970–979.

Nevo, S., Kadouri, N., and Abramson, J. (2019). Tuft cells: From the mucosa to the thymus. Immunol Lett 210, 1–9.

Parikh, K., Antanaviciute, A., Fawkner-Corbett, D., Jagielowicz, M., Aulicino, A., Lagerholm, C., Davis, S., Kinchen, J., Chen, H.H., Alham, N.K., et al. (2019). Colonic epithelial cell diversity in health and inflammatory bowel disease. Nature 567, 49–55.

Pelaseyed, T., Bergstrom, J.H., Gustafsson, J.K., Ermund, A., Birchenough, G.M., Schutte, A., van der Post, S., Svensson, F., Rodriguez-Pineiro, A.M., Nystrom, E.E., et al. (2014). The mucus and mucins of the goblet cells and enterocytes provide the first defense line of the gastrointestinal tract and interact with the immune system. Immunol Rev 260, 8–20.

Ringel, T., Frey, N., Ringnalda, F., Janjuha, S., Cherkaoui, S., Butz, S., Srivatsa, S., Pirkl, M., Russo, G., Villiger, L., et al. (2020). Genome-Scale CRISPR Screening in Human Intestinal Organoids Identifies Drivers of TGF-beta Resistance. Cell Stem Cell 26, 431–440 e438.

Sato, T., Stange, D.E., Ferrante, M., Vries, R.G., Van Es, J.H., Van den Brink, S., Van Houdt, W.J., Pronk, A., Van Gorp, J., Siersema, P.D., et al. (2011a). Long-term expansion of epithelial organoids from human colon, adenoma, adenocarcinoma, and Barrett’s epithelium. Gastroenterology 141, 1762–1772.

Sato, T., van Es, J.H., Snippert, H.J., Stange, D.E., Vries, R.G., van den Born, M., Barker, N., Shroyer, N.F., van de Wetering, M., and Clevers, H. (2011b). Paneth cells constitute the niche for Lgr5 stem cells in intestinal crypts. Nature 469, 415–418.

Sato, T., Vries, R.G., Snippert, H.J., van de Wetering, M., Barker, N., Stange, D.E., van Es, J.H., Abo, A., Kujala, P., Peters, P.J., et al. (2009). Single Lgr5 stem cells build crypt-villus structures in vitro without a mesenchymal niche. Nature 459, 262–265.

Schutgens, F., and Clevers, H. (2020). Human Organoids: Tools for Understanding Biology and Treating Diseases. Annu Rev Pathol 15, 211–234.

Smillie, C.S., Biton, M., Ordovas-Montanes, J., Sullivan, K.M., Burgin, G., Graham, D.B., Herbst, R.H., Rogel, N., Slyper, M., Waldman, J., et al. (2019). Intra- and Inter-cellular Rewiring of the Human Colon during Ulcerative Colitis. Cell 178, 714–730 e722.

Spohn, S.N., and Mawe, G.M. (2017). Non-conventional features of peripheral serotonin signalling - the gut and beyond. Nat Rev Gastroenterol Hepatol 14, 412–420.

Sugisawa, E., Takayama, Y., Takemura, N., Kondo, T., Hatakeyama, S., Kumagai, Y., Sunagawa, M., Tominaga, M., and Maruyama, K. (2020). RNA Sensing by Gut Piezo1 Is Essential for Systemic Serotonin Synthesis. Cell 182, 609–624 e621.

Vadgama, J.V., and Evered, D.F. (1992). Characteristics of L-citrulline transport across rat small intestine in vitro. Pediatr Res 32, 472–478.

van der Flier, L.G., van Gijn, M.E., Hatzis, P., Kujala, P., Haegebarth, A., Stange, D.E., Begthel, H., van den Born, M., Guryev, V., Oving, I., et al. (2009). Transcription factor achaete scute-like 2 controls intestinal stem cell fate. Cell 136, 903–912.

van der Vaart, J., and Clevers, H. (2020). Airway organoids as models of human disease. J Intern Med.

Wang, Y., Song, W., Wang, J., Wang, T., Xiong, X., Qi, Z., Fu, W., Yang, X., and Chen, Y.G. (2020). Single-cell transcriptome analysis reveals differential nutrient absorption functions in human intestine. J Exp Med 217.

Worthington, J.J., Reimann, F., and Gribble, F.M. (2018). Enteroendocrine cells-sensory sentinels of the intestinal environment and orchestrators of mucosal immunity. Mucosal Immunol 11, 3–20.

Yin, X., Farin, H.F., van Es, J.H., Clevers, H., Langer, R., and Karp, J.M. (2014). Niche-independent high-purity cultures of Lgr5+ intestinal stem cells and their progeny. Nat Methods 11, 106–112.

Zhou, J., Hegsted, M., McCutcheon, K.L., Keenan, M.J., Xi, X., Raggio, A.M., and Martin, R.J. (2006). Peptide YY and proglucagon mRNA expression patterns and regulation in the gut. Obesity (Silver Spring) 14, 683–689.

