## Supplemental Figures, Tables, and Methods for "Assessing donor-to-donor variability in human intestinal organoid cultures"

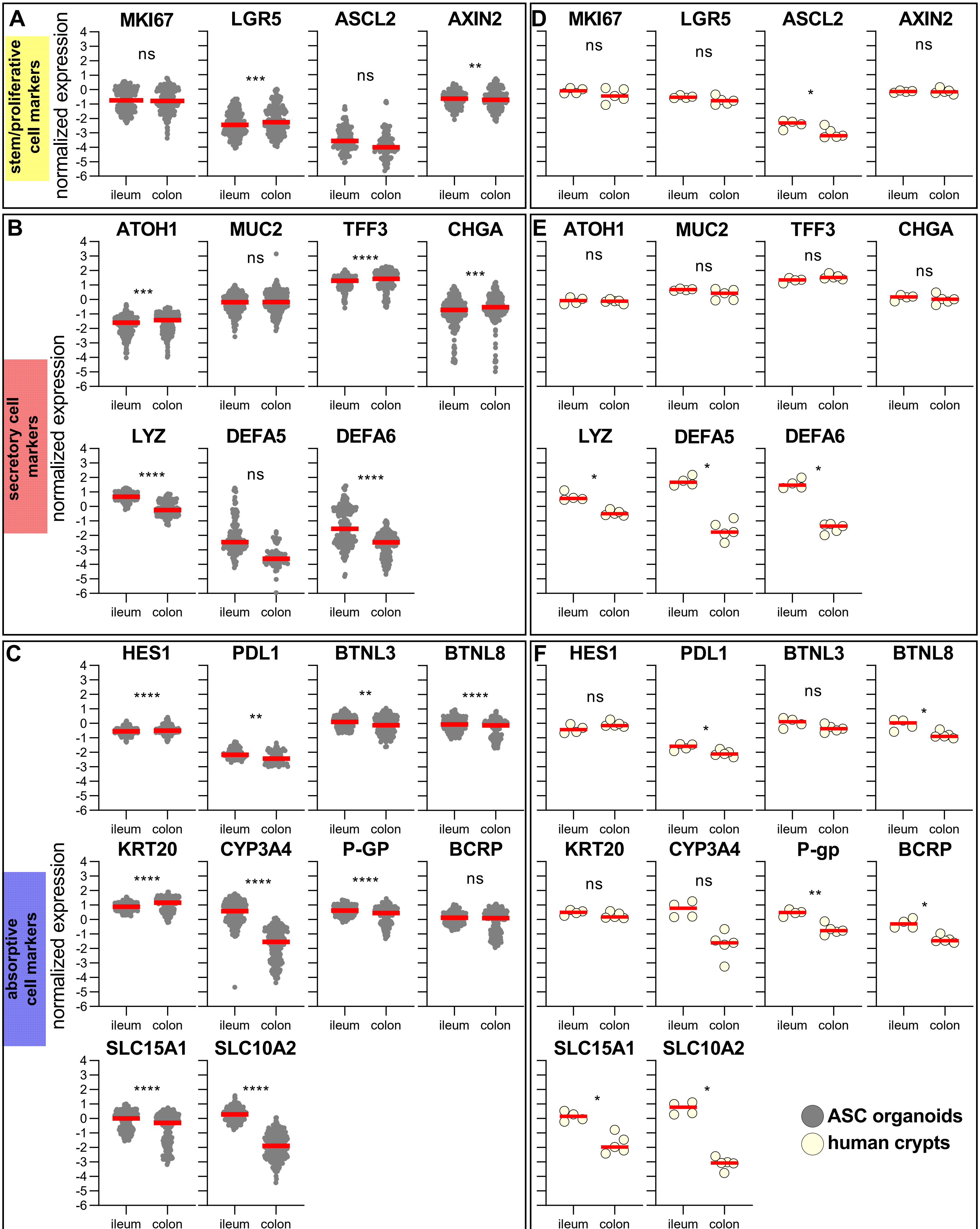
A
stem/proliferative cell markers
normalized expression
MKI67
LGR5
ASCL2
AXIN2
B
secretory cell markers
ATOH1
MUC2
TFF3
CHGA
LYZ
DEFA5
DEFA6
C
absorptive cell markers
HES1
PDL1
BTNL3
BTNL8
KRT20
CYP3A4
P-GP
BCRP
SLC15A1
SLC10A2
D
E
F
P-gp
ileum
colon
ns
ASC organoids
human crypts

Figure S2

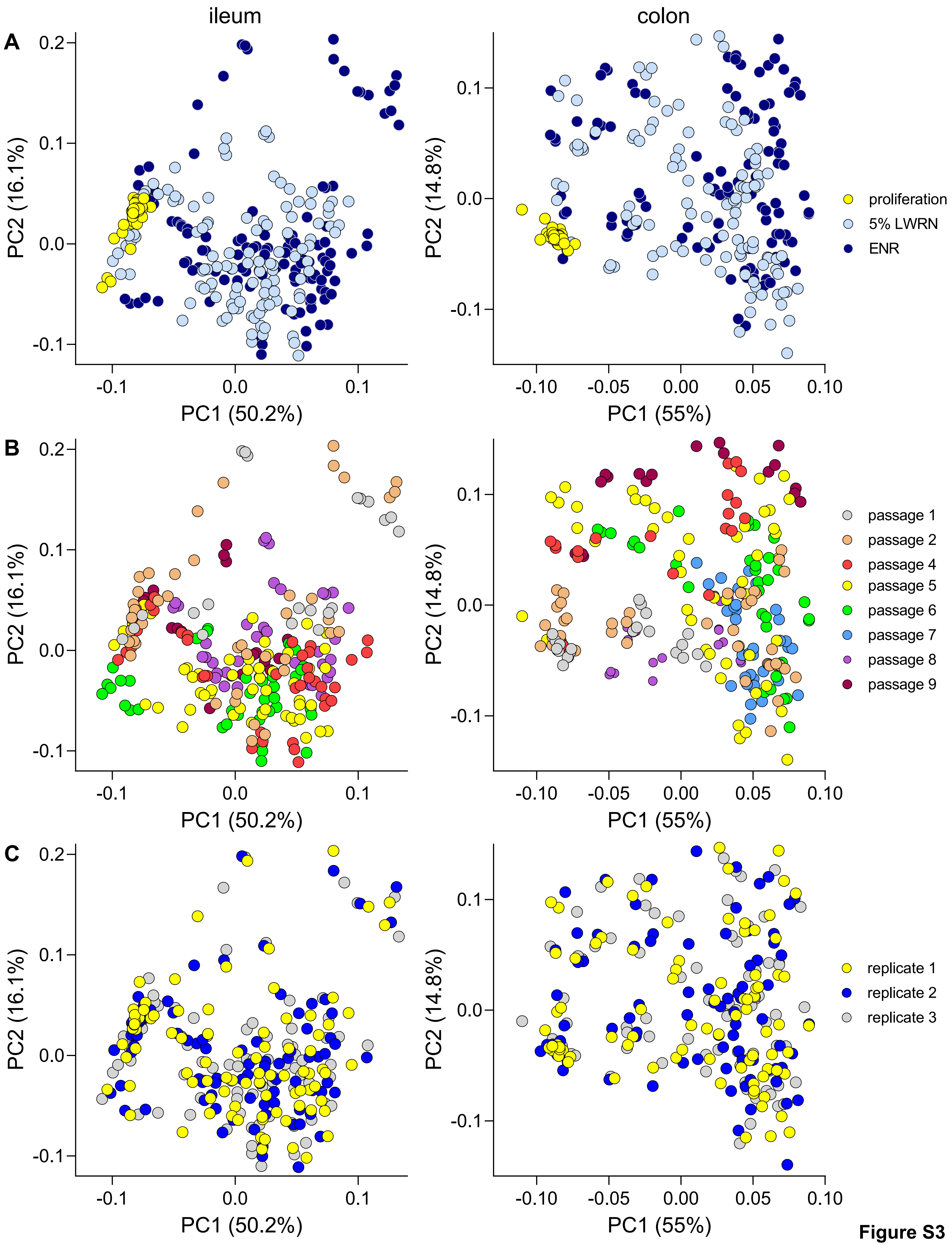

ileum
colon
A
B
C
PC1 (50.2%)
PC2 (16.1%)
PC1 (55%)
PC2 (14.8%)
proliferation
5% LWRN
ENR
passage 1
passage 2
passage 4
passage 5
passage 6
passage 7
passage 8
passage 9
replicate 1
replicate 2
replicate 3


**Figure S3**

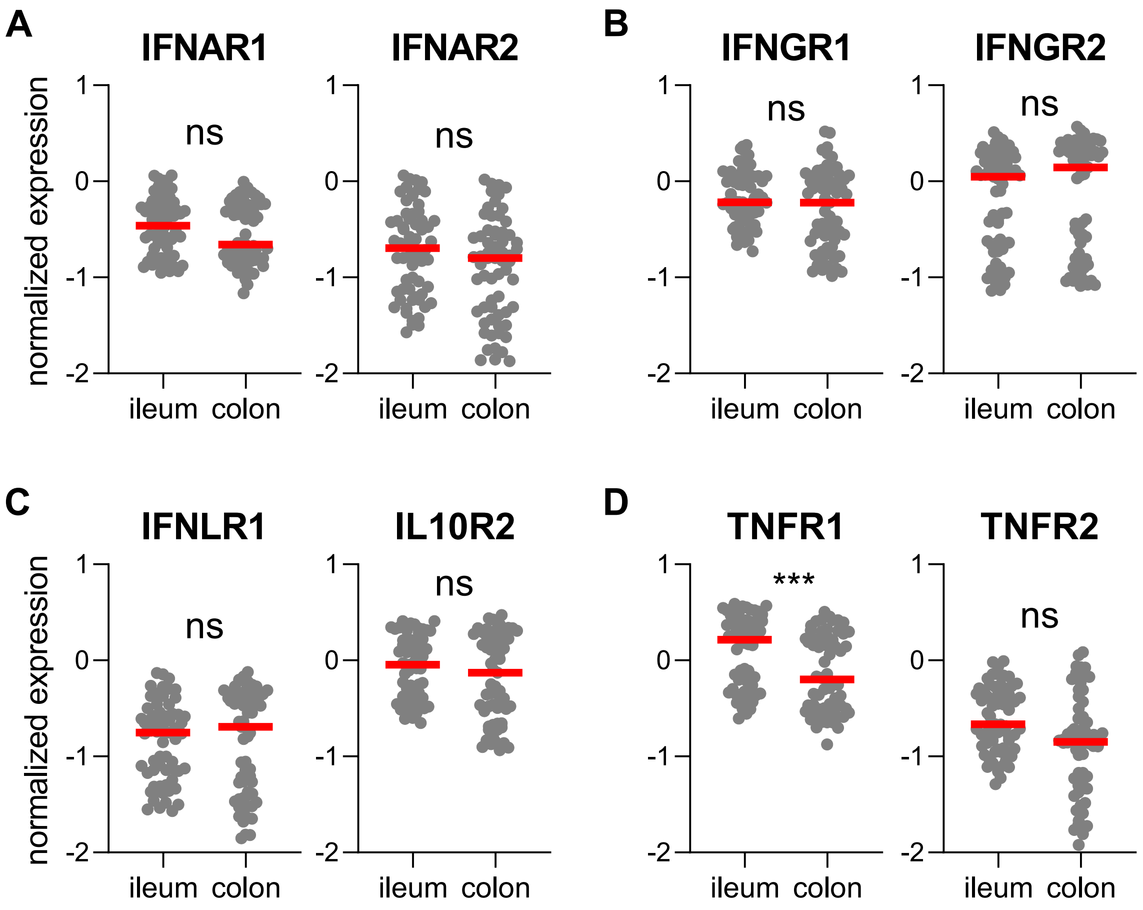
A
IFNAR1
IFNAR2
B
IFNGR1
IFNGR2
C
IFNLR1
IL10R2
D
TNFR1
TNFR2
normalized expression
ns
***
1
0
-1
-2
ileum
colon

**Figure S4**

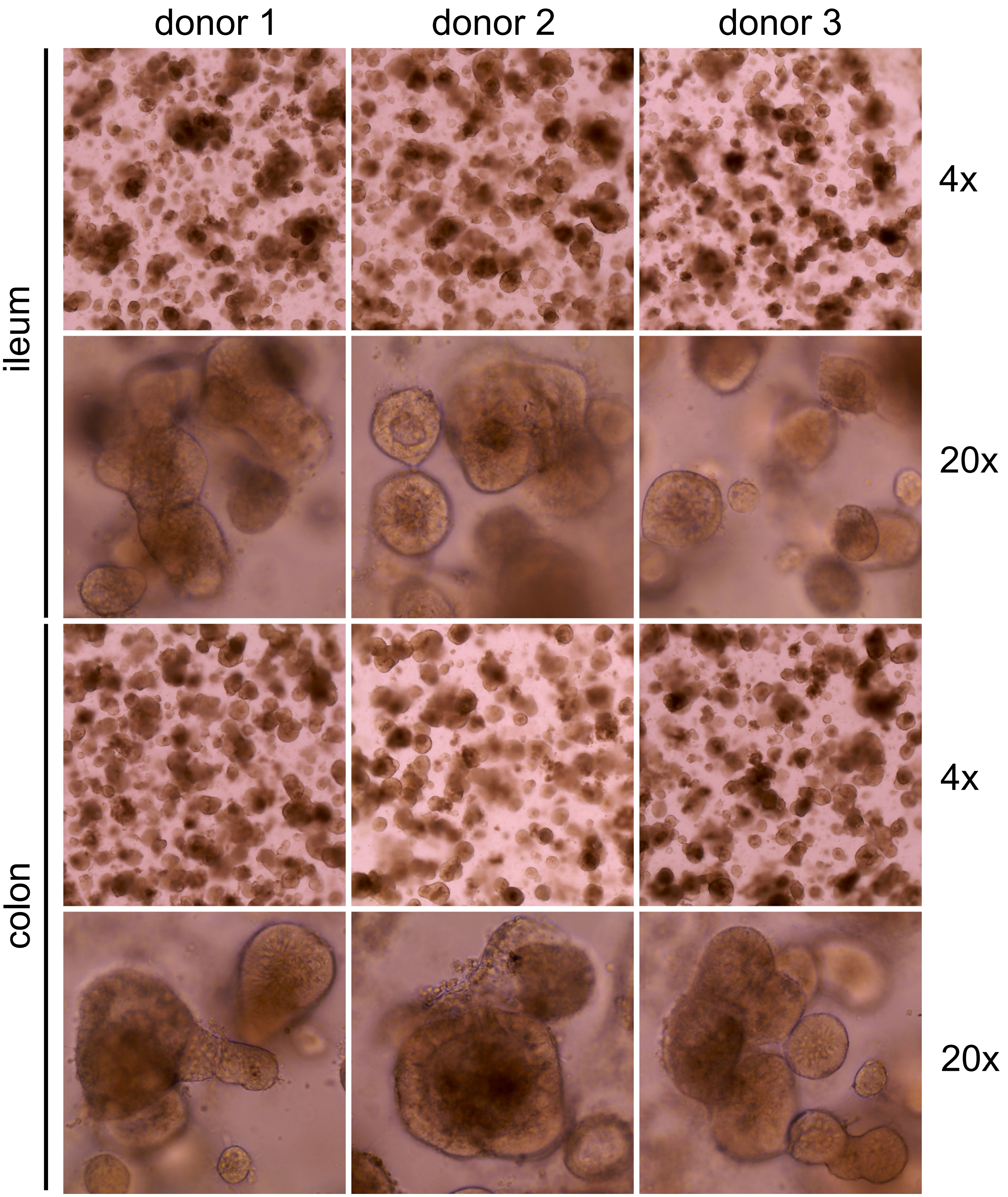


**Figure S5**

Supplemental Figure Legends:

Figure S2: Expression of markers in ileum or colon cultures
Expression of stem/proliferative (A and D), secretory (B and E), and absorptive (C and F) marker in both ileum and colon cultures (A-C) and human tissue (isolated crypts; D-F). Expression was quantified using qPCR and normalized to endogenous control (PPIB). In panels A-C, all samples (all time points, time courses) are included for each tissue type. Red line marks median value for each population. Symbols: ns, $p > 0.05$; * $p \leq 0.05$; ** $p \leq 0.01$; *** $p \leq 0.001$; **** $p < 0.0001$.

Figure S3: Assessing variability based on media, passage, replicate, and tissue type
Principal component analysis with samples marked by media type (A), passage number (B), or replicate number (C). Each symbol represents a separate RNA sample analyzed by qPCR.

Figure S4: Expression of immune receptors in ileum or colon cultures
Expression of type I (A), type II (B), and type III (C) interferon and TNF (D) receptors in both ileum and colon cultures (cumulative for all time points, time courses, and cultures). Expression, quantified by qPCR, was normalized to endogenous control (PPIB). Red line marks median value for each population. Symbols: ns, $p > 0.05$; * $p \leq 0.05$; ** $p \leq 0.01$; *** $p \leq 0.001$; **** $p < 0.0001$.

Figure S5: Brightfield analysis of ASC organoid cultures
Cultures from three donors were differentiated and on day 7, brightfield images were captured using a 4x (low magnification) and 20x (high magnification) objective lens.

**Supplementary Table 1**. LC/MS method for targeted analysis of central carbon metabolites

| LC Conditions | |
|---|---|
| Column | Agilent ZORBAX RRHD Extend-C18, 2.1 × 150 mm, 1.8 um (Part # 759700-902) |
| Guard column | ZORBAX Eclipse Plus C18, 2.1 mm, 1.8 µm, UHPLC guard column |
| Column temperature | 40 $^{0}$C |
| Needle wash | Methanol:water (50:50) with 15 mM glacial acetic acid |
| Mobile phase | (A) Water:Methanol (97:3) with 15 mM glacial acetic acid and 10 mM tributylamine (Acros Organics AC139320010)<br>(B) Methanol with 15 mM glacial acetic acid and 10 mM tributylamine<br>(D) Acetonitrile |
| Flow rate | Variable (see gradient table) |
| Gradient program | (see below) |
| Stop time | 40 mins |
| MS Ionization mode | ESI negative |

Gradient program:

| Time (min) | % A | % B | %D | Flow (mL/min) |
|---|---|---|---|---|
| 2.5 | 100 | 0 | 0 | 0.25 |
| 7.5 | 80 | 20 | 0 | 0.25 |
| 13.00 | 55 | 45 | 0 | 0.25 |
| 20.00 | 1 | 99 | 0 | 0.25 |
| 24.00 | 1 | 99 | 0 | 0.25 |
| 24.05 | 1 | 0 | 99 | 0.25 |
| 27.00 | 1 | 0 | 99 | 0.25 |
| 27.50 | 1 | 0 | 99 | 0.8 |
| 31.35 | 1 | 0 | 99 | 0.8 |
| 31.50 | 1 | 0 | 99 | 0.6 |
| 32.25 | 100 | 0 | 0 | 0.4 |
| 39.90 | 100 | 0 | 0 | 0.4 |
| 40.00 | 100 | 0 | 0 | 0.25 |

**Supplementary Table 2.** Metabolites identified by LC/MS QQQ analysis

| Compound Name | CAS ID | | Compound Name | CAS ID |
|---|---|---|---|---|
| 2-2-Dimethyl Succinic acid | 597-43-3 | | L-asparagine | 70-47-3 |
| 2-Deoxyadenosine 5-monophosphate | 653-63-4 | | L-Aspartic Acid | 56-84-8 |
| 2-Deoxyguanosine 5-diphosphate | 3493-09-2 | | L-Carnitine | 541-15-1 |
| 2-Deoxyguanosine 5-monophosphate | 902-04-5 | | L-Citrulline | 372-75-8 |
| 2-Deoxyribose 5-phosphate | 102916-66-5 | | L-Glutamic acid | 56-86-0 |
| 2-Deoxyuridine | 951-78-0 | | L-Glutamine | 56-85-9 |
| 2-Methyl-1-butanol | 1565-80-6 | | L-Gluthathione (oxidized) | 27025-41-8 |
| 2-Phosphoglyceric acid | 2553-59-5 | | L-Hydroxyglutaric acid | 13095-48-2 |
| 4-Hydroxybenzoic acid | 99-96-7 | | L-Isoleucine | 61-90-5 |
| 4-Hydroxy-L-glutamic acid | 2485-33-8 | | L-Kynurenine | 2922-83-0 |
| 4-Pyridoxic acid | 82-82-6 | | L-Leucine | 61-90-5 |
| 5-Hydroxy-3-indoleacetic acid | 54-16-0 | | L-Malic acid | 97-67-6 |
| Adenine | 73-24-5 | | L-Methionine | 63-68-3 |
| Adenosine | 58-61-7 | | L-Phenylalanine | 63-91-2 |
| Adenosine 3-5-cyclic monophosphate | 60-92-4 | | L-Proline | 147-85-3 |
| Adenosine 5-diphosphate | 58-64-0 | | L-Serine | 56-45-1 |
| Adenosine 5-monophosphate | 61-19-8 | | L-Sorbose | 87-79-6 |
| Adenosine 5-triphosphate | 56-65-5 | | L-Threonine | 72-19-5 |
| Adipic acid | 124-04-9 | | L-Tryptophan | 73-22-3 |
| Allantoin | 97-59-6 | | L-Tyrosine | 60-18-4 |
| alpha-D(+)Mannose 1-phosphate | 27251-84-9 | | Maleic acid | 110-16-7 |
| alpha-Ketoglutaric acid | 328-50-7 | | Malonic acid | 141-82-2 |
| Arabinose-5-phosphate | 13137-52-5 | | myo-Inositol | 87-89-8 |
| Argininosuccinic acid | 2387-71-5 | | N-Acetylglutamic acid | 5817-08-3 |
| beta-Nicotinamide adenine dinucleotide | 53-84-9 | | N-Acetylneuraminic acid | 131-48-6 |
| Cellobiose | 528-50-7 | | N-carbamoyl-DL-aspartic acid | 923-37-5 |
| Citramalic acid | 2306-22-1 | | N-Carbamyl-L-glutamic acid | 1188-38-1 |
| Citric acid | 77-92-9 | | O-Phosphorylethanolamine | 1071-23-4 |
| Creatine | 57-00-1 | | Orotic acid | 65-86-1 |
| Creatinine | 60-27-5 | | Phenylpyruvic acid | 156-06-9 |
| Cytidine-5-monophosphate | 63-37-6 | | Phosphoenolpyruvic acid | 138-08-9 |
| Deoxyguanosine 5-triphosphate | 2564-35-4 | | Pyridoxal hydrochloride | 65-22-5 |
| D-Gluconic acid | 526-95-4 | | Pyridoxine | 65-23-6 |
| Dihydroxyacetone phosphate | 57-04-5 | | Pyruvic acid | 127-17-3 |
| DL-2-Aminoadipic acid | 542-32-5 | | Riboflavin | 83-88-5 |
| D-Mannose | 31103-86-3 | | S-5-Adenosyl-L-homocysteine | 979-92-0 |
| D-pantothenic acid | 79-83-4 | | Salicylic acid | 69-72-7 |

| | | | | |
|---|---|---|---|---|
| D-Sedoheptulose-7-phosphate | 2646-35-7 | | Succinic acid | 110-15-6 |
| D-Xylose | 58-86-6 | | Taurine | 107-35-7 |
| Flavin adenine dinucleotide | 146-14-5 | | Taurocholic acid | 83830-80-2 |
| Galactonic acid | 576-36-3 | | Thiamine | 67-03-8 |
| gamma-Glu-Cys | 636-58-8 | | Thymine | 65-71-4 |
| Glyceric acid | 473-81-4 | | trans-4-Hydroxy-L-proline | 51-35-4 |
| Guanosine | 118-00-3 | | Uracil | 66-22-8 |
| Guanosine 5-triphosphate | 86-01-1 | | Uric acid | 69-93-2 |
| Hypoxanthine | 68-94-0 | | Uridine | 58-96-8 |
| Inosine | 58-63-9 | | Uridine 5-diphosphate | 58-98-0 |
| Inosine 5-triphosphate | 132-06-9 | | Uridine 5′-diphosphogalactose | 2956-16-3 |
| Isopentyl acetate | 123-92-2 | | Uridine 5-diphosphoglucose | 133-89-1 |
| Itaconic acid | 97-65-4 | | Uridine 5-monophosphate | 58-97-9 |
| Ketovaleric acid | 1821-02-9 | | Uridine 5-triphosphate | 63-39-8 |
| Lactic acid | 50-21-5 | | Xanthine | 69-89-6 |
| L-Arabinose | 5328-37-0 | | Xanthosine | 146-80-5 |
| L-Arabitol | 7643-75-6 | | Xylitol | 87-99-0 |

Supplemental Experimental Procedures:

*Human ASC organoid culture*

Cultures were derived from donor tissue, essentially as previously described (Miyoshi and Stappenbeck, 2013; Sato et al., 2011a). Donor information is summarized in Table 1. Briefly, small intestine and colon tissue were cut open longitudinally and rinsed with cold PBS containing gentamicin and amphotericin B (Invitrogen R01510) to remove debris. Mucosa was dissected away from muscle tissue using fine scissors and forceps. Mucosa was sliced into $1mm^2$ pieces using a tissue chopper. Tissue pieces were incubated in 10mM DTT at room temperature twice for 10 minutes with gentle agitation. DTT was replaced with ice cold 5mM EDTA and tissue was incubated at 4ºC with gentle agitation. Crypts were released by shaking tubes by hand and filtering through a 100µm strainer. Isolated crypts were embedded in Matrigel (Corning 356237), plated in 24 well dishes, and overlaid with growth media as described previously (Sato et al., 2011a). Additionally, an aliquot of isolated crypts was lysed for RNA extraction (except for donor 3 ileum) and used as reference for original tissue gene expression analysis. After first passage, ASC organoids were cultured in 50% LWRN conditioned media containing 10µM Y27632 and 10µM SB431542 (Miyoshi and Stappenbeck, 2013). Media was changed every 2-3 days and cultures were passaged once per week.

*Differentiation time course cultures*

ASC organoid cultures were differentiated as follows. ASC organoids were seeded in 24 well dishes and fed with proliferation media (50% LWRN + Y27632 + SB431542) as described previously (Miyoshi and Stappenbeck, 2013). This media is composed of conditioned media containing Wnt3a, Rspo3, and noggin along with Rho-associated protein kinase (ROCK) and transforming growth factor (TGF) β type I receptor inhibitors. This complex media does not contain EGF, previously shown to be critical for ASC organoid cultures (Sato et al., 2011a). It is thought that this activity is provided by undefined components present in serum. Similarly, it has been shown that noggin could be eliminated from this system without negatively affecting ASC organoid growth (Miyoshi and Stappenbeck, 2013). Media was changed on day 2 post seeding. Three days after seeding in 24 well dishes, media on cultures was changed to either 5% LWRN conditioned media (no Y27632 or SB431542) or a defined media containing 50ng/mL EGF, 100ng/mL Noggin, and 250ng/mL Rspo1 in Advanced DMEM/F12 (referred to as ENR media). ENR media was composed of purified EGF, Noggin, and Rspondin-1 added to base media. ENR media was used as it is composed entirely of defined components and is serum-free. 5% LWRN media was evaluated because of ease of preparation and low cost but keeping in mind the potential liabilities posed by presence of serum. Additional differences between the two media formulations is presence of Wnt. 5% LWRN media contains low amount of Wnt3a, while ENR media contains no added Wnt. As Wnt signaling is critical for stem cell maintenance in the intestine (Krausova and Korinek, 2014), 5% LWRN media could potentially stimulate low-level stem cell maintenance during the differentiation process. Low levels of Wnt3a in media could replenish the culture as cells terminally differentiate. However, this low level Wnt3a could skew differentiation patterns. As an example, as Paneth cell development and maintenance required Wnt signaling (Gassler, 2017; Ireland et al., 2005; Sato et al., 2011b), more Paneth cells may be found in cultures differentiated in 5% LWRN media. Because of both technical and potential impact on signaling pathways, both media formulations were included.

Media was changed on days 5, 7, and 9. Samples were collected for RNA purification on days 2, 4, 7, and 10. For RNA purification, 350µL RLT + 2-ME (Qiagen RNeasy kit) was added to each well. Lysates were stored at -80ºC until purification. The number of time course carried out for each culture is as follows: 3, donor 1 ileum; 3, donor 1 colon; 3, donor 2 ileum; 3, donor 2 colon; 2, donor 3 ileum; 3, donor 3 colon; 3, donor 4 ileum; 2, donor 5, colon; 2, donor 6 ileum; 2, donor 6 colon. For each time course, 3 cell culture replicates (separate wells processed independently) were included for every condition. Three donor cultures (both ileum and colon cultures) were routinely maintained. For assays which used fewer than all six donors (Figures 4-7), no attempt was made to select best performing or most responsive cultures.

*Quantitative PCR*

RNA was purified from ASC organoid cultures using RNeasy kit (Qiagen 74104), always including on-column DNAse digest (Qiagen RNase-Free DNase Cat # 79254). cDNA was generated using SuperScript IV (Invitrogen 11756500) and diluted 5x with water to generate PCR template. TaqMan Fast Master Mix (Invitrogen 4444557) and TaqMan assays (Invitrogen) were used to assess gene expression and were used as indicated by the manufacturer. Quantitative PCR was carried out on a QuantStudio 12K Flex instrument (Applied Biosystems). Data were normalized to an endogenous control (PPIB) using the ΔCt method as previously described (Schmittgen 2008). All markers are summarized in Table 2.

*Principal component and correlation analysis*

Principal component analysis (PCA) was performed in the R statistical computing environment (Team, 2019) by singular value decomposition (SVD). Data was first scaled and centered prior to SVD. SVD was performed with the function SVDmiss in the SpatioTemporal package (Lindstrom et al., 2019). This function iteratively performs SVD while replacing missing values by linear regression of the columns onto the first four SVD components. Coordinates for the first two principle components were then plotted using Graphpad Prism. Coloring indicates membership in groups of interest such as day or donor. Total percent variation accounted for by each PC is given in the axes' labels. Correlations were performed in the R statistical computing environment . Samples were subset to those containing only the 5% LWRN and then further subset to those from only the day 7 timepoint. Correlations were performed with the cor function in R using the Pearson correlation metric and missing data was handled by casewise deletion using the complete.obs setting for the use argument.

*Immunofluorescence staining*

ASC organoids were removed from Matrigel using cell recovery solution (Corning 354253) by incubating on ice for 30min (with occasional inversion to mix) and fixed using 4% paraformaldehyde in 1X PBS for 60min at ambient temperature. Fixed ASC organoids were washed with TBS containing 0.5% BSA three times and permeabilized for 2hrs at ambient temperature in TBS containing 0.5% BSA and 1% Triton X-100. ASC organoids were blocked in TBS containing 5% BSA and 0.1% Triton X-100 overnight at 4ºC. Incubation with primary antibodies were carried out at 4ºC overnight. Antibodies used were: MUC2 (1:200; Dako M731329-2), Lyzosyme (1:500; Invitrogen PA5-16668), Chromogranin A (1:100; Abcam ab15160), E-cadherin (1:100; BD 610181), or E-cadherin (1:200 Cell Signaling Technology 3195S). Secondary antibodies were all from Invitrogen and were used at 1:250 dilution. Actin and DNA were detected with phalloidin (1:400; Invitrogen A22287) and Hoechst 33342 (1:2000;

Invitrogen H3570) respectively. Stained ASC organoids were mounted on slides using ProLong Gold mounting media (Invitrogen P36934) and imaged on a Zeiss LSM800 confocal microscope using a Plan-Apochromat 63x/1.40NA lens. Contrast was adjusted linearly in Zeiss Zen software for each channel and merged 8-bit tiff images were exported. Figure was assembled in Adobe Photoshop. For quantification, multiple images were captured from each marker and culture. For each culture hundreds of cells were analyzed: >500 cells for all samples and markers except for lysozyme staining in donor 2 (307 cells) and donor 3 (473 cells) ileum cultures. DNA stain (Hoechst) was used to identify all cells. Cell type markers were used to identify specific differentiated cells. A ratio of cell type marker to total cells was calculated and presented as "% of cells".

*Hormone secretion assay*

ASC organoids were plated as described for time course experiments (in 24 well dishes), changing to differentiation media (5% LWRN CM) on day 3 after plating. On day 7 after plating ASC organoids were washed 3x with 500µL PBS (+ Calcium/Magnesium) + 10mL HEPES. Hormone secretion was stimulated as previously described (Billing et al., 2018) by adding 200µL per well PBS (+ Calcium/Magnesium) + 10mL HEPES + 10µM forskolin (Sigma F6886) + 10µM IBMX (Sigma I5879) + 10mM glucose. Plates were incubated at 37ºC/5% $CO_2$ overnight and supernatant collected. Four wells were collected for each condition. Commercially-available ELISA kits were used to quantify hormone levels (PYY: Millipore EZHPYYT66K; GPL1: Millipore EZGLP1T-36K; Serotonin: Enzo ADI-900-175).

*Metabolomics analysis*

Samples were analyzed by targeted LC/MS analysis, using and Agilent 6470 Triple Quadrupole (QQQ) mass spectrometer, in negative ionization mode, coupled to an Agilent 1290 Infinity II HPLC with quaternary pump. Metabolites were separated using an Agilent ZORBAX RRHD Extend-C18 (2.1 × 150 mm, 1.8 µm) column with the following mobile phases: (A) H2O:methanol (97:3) with 15 mM glacial acetic acid and 10 mM tributylamine; (B) methanol with 15 mM glacial acetic acid and 10 mM tributylamine; (D) acetonitrile. Multiple Reaction Monitoring (MRM) transitions for the central carbon metabolites were from the Agilent Metabolomics MRM Database and Method. Data was analyzed using Agilent Quantitative Analysis B.08 and Mass Profiler Professional version 14.9.1 software. Additional method information is in Supplementary Tables 1 and 2.

For each sample, media was aspirated, and ASC organoids and Matrigel were washed with PBS. Cell recovery solution (Corning # 354253) was added to Matrigel and transferred to a homogenization tube (Fisher #15-340-153), which was centrifuged at 500xg at 4 ºC for 5 minutes. The supernatant was aspirated, and 500 µL of ice cold extraction solvent (acetonitrile: methanol: water; 40:40:20) was added to each tube and homogenized with a bead mill homogenizer (Omni Bead Ruptor 19-040E) for 30 seconds at -5 ºC at 5.0 m/s. ASC organoids were allowed to extract for 10 minutes at -20 °C. After 10 minutes, the tubes were centrifuged for 5 minutes at 4 ºC at 14,000 x g. The supernatant (350 µL) was transferred to a clean 1.5 mL microcentrifuge tube and dried under nitrogen. Each sample was dissolved in 30 µL water:methanol (80:20) for LC/MS analysis.

*Statistics*

Comparisons were performed using a two-tailed Student's t test, calculated in either Microsoft Excel or Graphpad Prism. Asterisks indicate p value ranges as follows: ns, $p > 0.05$; * $p \leq 0.05$; ** $p \leq 0.01$; *** $p \leq 0.001$; **** $p < 0.0001$.

References:

Billing, L.J., Smith, C.A., Larraufie, P., Goldspink, D.A., Galvin, S., Kay, R.G., Howe, J.D., Walker, R., Pruna, M., Glass, L., *et al.* (2018). Co-storage and release of insulin-like peptide-5, glucagon-like peptide-1 and peptideYY from murine and human colonic enteroendocrine cells. Mol Metab *16*, 65-75.
Gassler, N. (2017). Paneth cells in intestinal physiology and pathophysiology. World J Gastrointest Pathophysiol *8*, 150-160.
Ireland, H., Houghton, C., Howard, L., and Winton, D.J. (2005). Cellular inheritance of a Cre-activated reporter gene to determine Paneth cell longevity in the murine small intestine. Dev Dyn *233*, 1332-1336.
Krausova, M., and Korinek, V. (2014). Wnt signaling in adult intestinal stem cells and cancer. Cell Signal *26*, 570-579.
Lindstrom, J., Szpiro, A., Sampson, P.D., Bergen, S., and Oron, A.P. (2019). SpatioTemporal: Spatio-Temporal Model Estimation. R package version 1191.
Miyoshi, H., and Stappenbeck, T.S. (2013). In vitro expansion and genetic modification of gastrointestinal stem cells in spheroid culture. Nat Protoc *8*, 2471-2482.
Sato, T., Stange, D.E., Ferrante, M., Vries, R.G., Van Es, J.H., Van den Brink, S., Van Houdt, W.J., Pronk, A., Van Gorp, J., Siersema, P.D., *et al.* (2011a). Long-term expansion of epithelial organoids from human colon, adenoma, adenocarcinoma, and Barrett's epithelium. Gastroenterology *141*, 1762-1772.
Sato, T., van Es, J.H., Snippert, H.J., Stange, D.E., Vries, R.G., van den Born, M., Barker, N., Shroyer, N.F., van de Wetering, M., and Clevers, H. (2011b). Paneth cells constitute the niche for Lgr5 stem cells in intestinal crypts. Nature *469*, 415-418.
Team, R.C. (2019). R: A language and environment for statistical computing. R Foundation for Statistical Computing.
